# YB-1 IS REQUIRED FOR THE GENESIS AND METASTATIC CAPACITY OF HUMAN BREAST CANCER

**DOI:** 10.1101/372524

**Authors:** Sylvain Lefort, Amal El-Naggar, Susanna Tan, Shane Colborne, Gian Luca Negri, Davide Pellacani, Martin Hirst, Barry Gusterson, Gregg B. Morin, Poul H. Sorensen, Connie J. Eaves

## Abstract

Breast cancer heterogeneity has made it challenging to elucidate shared mechanisms that underpin properties that are critical to their growth *in vivo*. Here, we interrogate the role of YB-1 protein in the *in vivo* tumorigenic activity of *de novo* well as cell line models of human breast cancer. Short-hairpin RNA-mediated knockdown of YB-1 in MDA-MB-231 cells blocked both their local tumour-forming and lung-colonizing activity in transplanted immunodeficient mice. YB-1 knockdown also revealed its important role at early stages of human mammary cell transformation in the generation of invasive ductal carcinoma and ductal carcinoma *in situ* produced in mice transplanted with freshly isolated human mammary cells transduced, respectively, with *KRAS^G12D^*, or myristoylated-AKT1. Conversely, upregulated expression of YB-1 in the poorly tumorigenic T47D cells enhanced this activity. Mechanistically, reducing YB-1 levels in MDA-MB-231 cells impaired their induction of HIF1α, and G3BP1, known YB-1 translational targets and key elements of a stress-adaptive program.

## INTRODUCTION

Mammalian Y-box binding protein-1 (YB-1) is a member of the family of DNA/RNA binding proteins with a conserved cold-shock domain (CSD). Mammalian CSD proteins are widely expressed and involved in many fundamental processes including DNA repair as well as mRNA transcription, splicing, stabilization, and translation(1, 2). In many tumour types, most notably those with metastatic activity, elevated YB-1 expression correlates with drug resistance and poor survival (3, 4). In human sarcoma cells, we previously showed that YB-1 can drive metastasis through its ability to bind directly to *HIF1A* (5) and *NFE2L2* (6) mRNAs and thereby enhance their translation in cells responding to diverse stress conditions. YB-1 has also been shown to promote stress-induced stress granule (SG) formation in pancreatic and colon cells (7), and we have demonstrated that this involves the translational activation of *G3BP1* mRNAs in multiple human tumour cell types (8). We have also previously shown that YB-1 can contribute to the acquisition of a stress-related increase in invasive and metastatic properties of human malignant cells (9). However, the role of YB-1 in the initial stages of malignant transformation of human cells in an *in vivo* context remains poorly defined, due both to the paucity of primary samples of viable tissue at this stage and of experimental models of *de novo* human cancer development *in vivo* from cells isolated directly from normal human tissue.

To overcome this constraint, we have taken advantage of a system we have described recently that allows invasive ductal carcinomas (IDCs) to be reproducibly and rapidly obtained from freshly isolated normal human mammary cells transduced with a lentiviral vector encoding a *KRAS^G12D^* cDNA (10, 11). This approach efficiently transforms cells from two of the three major subsets of human mammary cells, specifically the two that proliferate in response to epidermal growth factor (EGF). These two cell types are referred to as basal cells (BCs) and luminal progenitors (LPs), the latter representing a phenotypically and biologically distinct subset of the luminal cell compartment (12). Tumours produced when several thousand *KRAS^G12D^*–transduced BCs or LPs are transplanted into immunodeficient mice are highly polyclonal and phenotypically heterogeneous, with a variable content of cells positive for estrogen receptor (ER), progesterone receptor (PR), human epidermal growth factor receptor-2 (HER2), EGFR, Ki67 and cytokeratins (CK) 8/18 expression (10). Interestingly, we found that neither mutant *TP53* or *PIK3CA* cDNAs alone had detectable oncogenic activity in either the BCs or LPs, and neither enhanced that obtained with *KRAS^G12D^* alone (10).

Here we demonstrate a shared *in vivo* growth dependence on YB-1 of transformed human mammary cells that span a broad spectrum of tumorigenic activity in transplanted immunodeficient mice. The models investigated include a new model of ductal carcinoma *in situ* (DCIS) that we now show is produced *de novo* from normal human mammary cells transduced with a vector encoding a myristoylated form of *AKT1* (*myr-AKT*), as well as the cells of *de novo KRAS^G12D^*-induced IDCs, and two human breast cancer cell lines with different *in vivo* tumour-initiating and metastatic activities. Importantly, the pervasive role of YB-1 includes an activation of known mediators of a broad cytoprotective stress response.

## RESULTS

### Identification of an association of altered KRAS and AKT activity with increased YB-1 expression, and activated stress responses in patients’ breast cancers

Examination of 3 large breast cancer datasets (13–15) showed that elevated levels of *YBX1* (hereafter referred to as *YB-1*) transcripts were associated with a reduced overall survival of patients with ER- breast cancers, most notably in those with metastatic disease. Elevated expression of YB-1 was also positively associated with a gain of function or amplification of the *KRAS, ERBB2* and *PIK3CA* genes, or deletions of the *TP53* gene, but not with amplified *HRAS* or *NRAS* mutations (Fig 1A-C and Appendix Fig S1A-D and data not shown). Previous studies have indicated that YB-1 can alter the translational control in malignant cells of a large number of proteins involved in cytoprotective responses to stresses they encounter, including HIF1α, G3BP1, and NRF2 (5, 6, 8, 16). From analyses of the same 3 patient breast cancer datasets, we found that *HIF1A* transcripts and its transcriptional targets, *CAIX* and *VEGFA* (17) and *G3BP1* (but not *NRF2*) were all significantly higher in breast cancers that exhibit gain of function or amplified *KRAS* (Fig 1D and E and Appendix Fig S2A-C). Elevated levels of *HIF1A*, its target *CAIX*, and *VEGFA* transcripts were also noted in tumours with a gain of function or amplification of *YB-1*, although this was not the case for *G3BP1* or *NFE2L2* (encoding NRF2; Appendix Fig S2D). Notably, analyses of these datasets revealed that *YB-1, HIF1A, CAIX* and *G3BP1* transcript levels were also increased in tumours with amplified *AKT1* and/or increased *AKT1* expression (Fig 1F and G, and Appendix Fig S2E-H). Together, these data suggest that induction of a YB-1-mediated stress response pathway might play a role in supporting the malignant properties of human breast cancers *in vivo*

**Figure 1.**
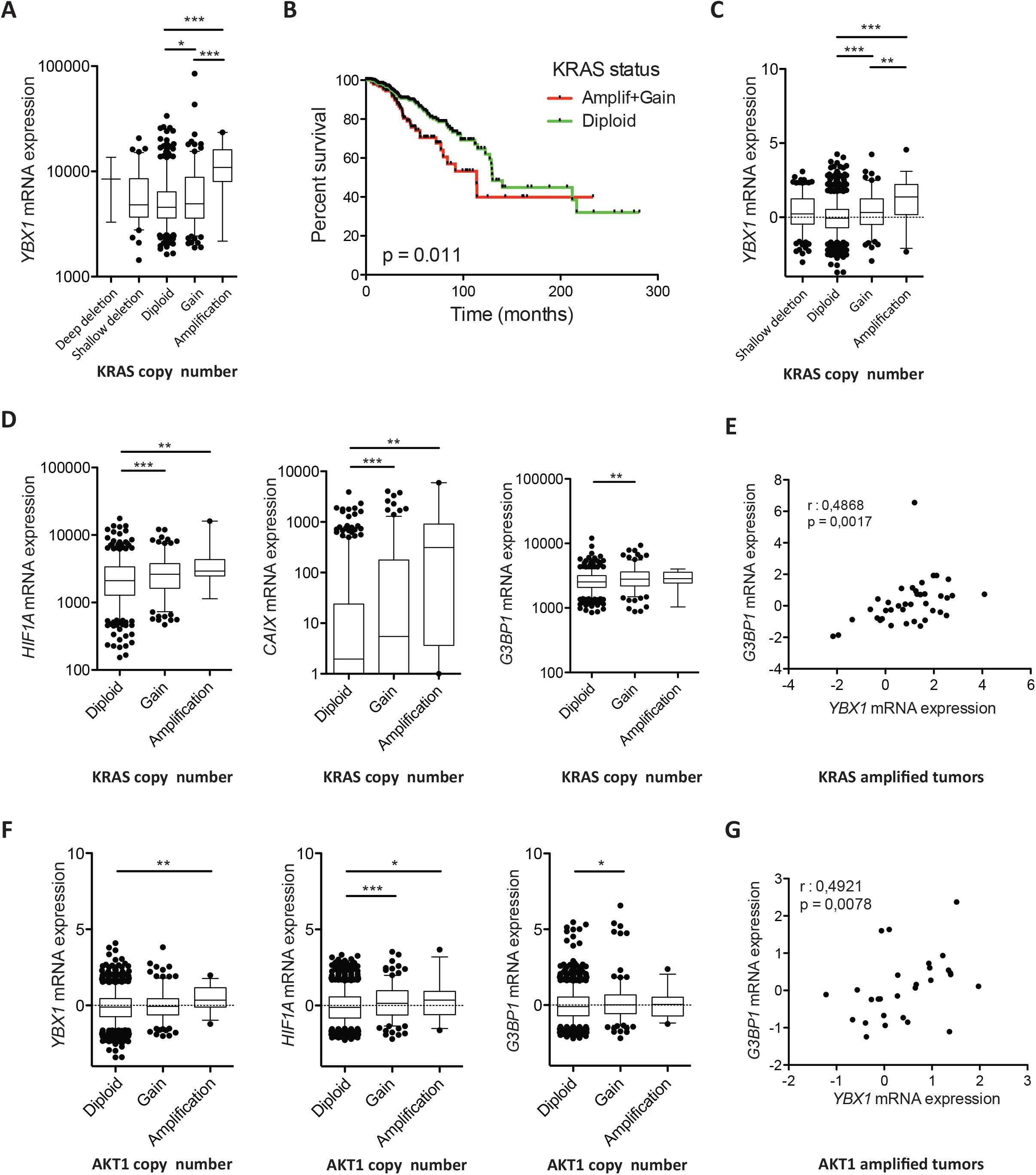
Patients’ *KRAS*-amplified tumours express high levels of *YB-1* and associated stress response genes. **A.** *YB-1* mRNA levels compared to *KRAS* copy number status in invasive breast carcinoma samples in the TCGA dataset. Values for *YB-1* are shown as RPKMs. **B.** Kaplan-Meier curves of overall survival (OS) for the TCGA cohort, with respect to *KRAS* copy number (N=206 for tumours with amplified *KRAS* or a gain of function *KRAS* gene, and N=522 for tumours with diploid *KRAS*). **C.** *YB-1* mRNA levels compared to *KRAS* copy number status in invasive breast carcinoma samples in the METABRIC dataset. Values for *YB-1* are shown as RPKMs. **D.** *HIF1A* (left panel) *CAIX* (middle panel) and *G3BP1* (right panel) mRNA expression according to *KRAS* copy number status in invasive breast carcinomas in TCGA dataset. Values for *HIF1A, CAIX and G3BP1* are shown as RPKMs. **E.** Scatter plot of *YB-1* and *G3BP1* mRNA expression in amplified-*KRAS* invasive breast carcinomas. **F***YB-1* (left panel) *HIF1A* (middle panel) and *G3BP1* (right panel) mRNA expression according to *AKT1* copy number status in invasive breast carcinomas in METABRIC dataset. Values for *YB-1*, *HIF1A and CAIX* are shown as RPKMs. **G.** Scatter plot of *YBX1* and *G3BP1* mRNA expression in amplified-*AKT1* invasive breast carcinomas. P-values in panels (A, C-D) were determined by Student’s t-test, \**P*<0.05, \*\**P*<0.01, \*\*\**P*<0.001.

### YB-1 is required for the tumorigenic and metastatic activities of MDA-MB-231 cells in transplanted immunodeficient mice

As a first approach to examining the biological significance of elevated YB-1 expression on the *in vivo* tumorigenic properties of human breast cancer cells with altered *KRAS* or *AKT* expression, we first focused on determining the effect of reducing YB-1 protein levels in xenografts of MDA-MB-231 cells. This human breast cancer cell line contains a *KRAS^G13D^* mutation (18), displays elevated YB-1 expression (19) and displays aggressive tumorigenic and metastatic activities when transplanted into highly immunodeficient female mice (11, 20). Transduction of these cells with lentiviral vectors encoding short hairpin (*sh*)*YB-1* constructs reduced their YB-1 protein content by 90% compared to controls (Fig 2A, Appendix Fig S3A-B). Reduced YB-1 expression, in turn, caused a marked decrease in both the size (Fig 2B and Appendix Fig S3C, D) and YB-1 content (Appendix Fig S3E) of the tumours produced at their subcutaneous (SQ) sites of injection. Parallel experiments in which mice were injected intravenously with the same test or control shRNA-transduced cells showed dramatically reduced metastatic activity of the *shYB-1*-transduced cells in the lungs as compared to control cells (Fig 2C-D and Appendix Fig. S3F-H). Taken together, these results demonstrate the strong YB-1 dependence of both the *in situ* and metastatic growth of MDA-MB-231 cells in xenografted immunodeficient mice.

**Figure 2.**
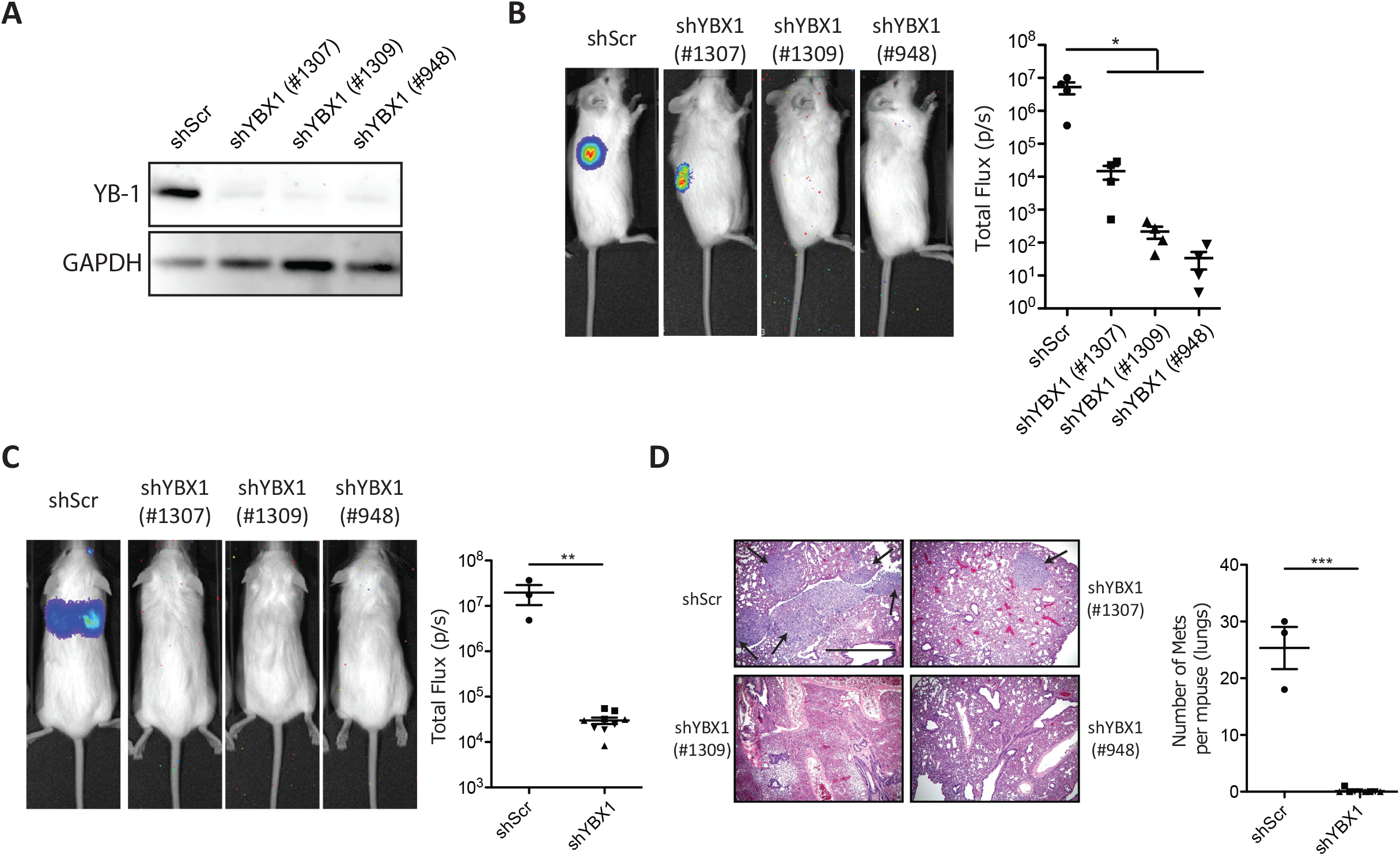
Suppressed YB-1 expression impairs the generation of local and metastatic tumour formation by the MDA-MB-231 cells. **A.** Western blots of MDA-MB-231 cells transduced with *shYB-1(#1307), shYB-1(#1309)* or *sh*Y*B-1(#948)* or a control *Scr* vector, showing YB-1 protein levels (relative to GAPDH). **B.** Representative pictures of bioluminescence signals in mice injected SQ with MDA-MB-231 cells transduced with *shYB-1(#1307), shYB-1(#1309), shYB-1(#948)* or *shScr.* Dot plots show the bioluminescence exhibited by these tumours 34 days post-transplant. **C.** Representative pictures of bioluminescence signals obtained in mice 34 days after being injected intravenously with MDA-MB-231 cells transduced with *shYB-1* or *shScr* vectors. Dot plot shows the measured bioluminescence exhibited by the progeny of these cells generated *in vivo*. **D.** Representative photomicrographs of H&E- or YB-1-stained sections of lungs of mice injected intravenously with MDA-MB-231 cells transduced with *shYB-1* or *shScr* vectors. Dot plot shows the number of metastasis per mouse (2 lungs). P-values in panels (B-D) were determined by Student’s t-test, \**P*<0.05, \*\**P*<0.01, \*\*\**P*<0.001.

### Loss of MDA-MB-231 cell tumorigenic activity by YB-1 inactivation is associated with evidence of a suppressed stress response

Despite the marked effects of reduced YB-1 levels on the growth of the *shYB-1*-transduced MDA-MB-231 cells *in vivo*, cell-cycle distribution analysis did not reveal any alteration of their proliferative activity in standard 2D cultures (Fig 3A), consistent with previous reports (5). However, even a 24-hour incubation under non-adherent culture conditions that promote the formation of spheroids (6), resulted in a blocked expression of HIF1α, CAIX, G3BP1, and NRF2 in the *shYB-1*-transduced cells in comparison to similarly cultured control-transduced cells (Fig 3B). HIF1α expression was also reduced in *shYB-1*-transduced MDA-MB-231 cells cultured under hypoxic conditions in comparison to their control vector-transduced counterparts (Fig 3C).

**Figure 3.**
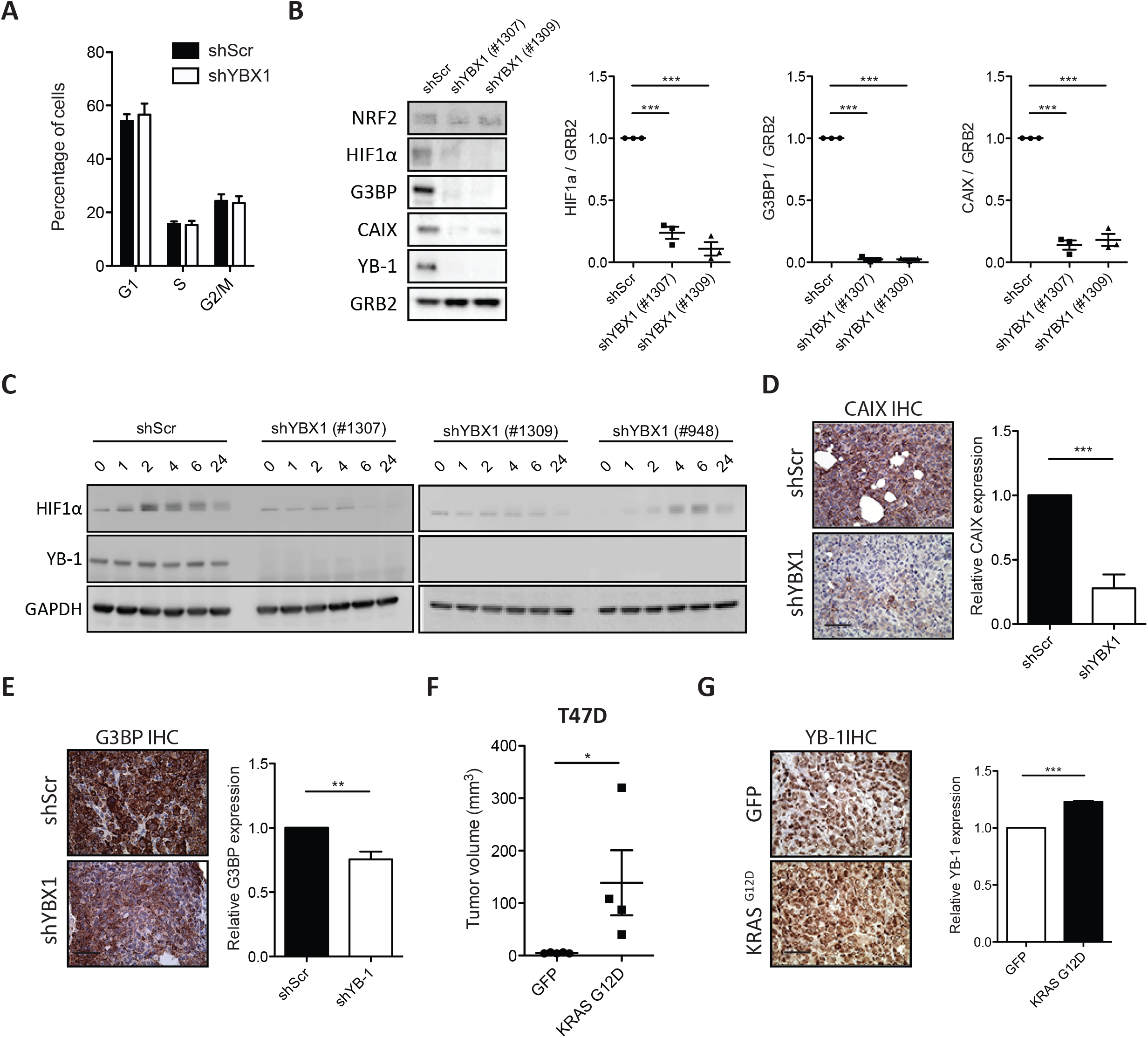
Loss of tumorigenic activity by YB-1 inactivation is associated with a suppression of stress response proteins. **A.** Cell cycle analysis of *shYB-1*- or *shScr*-transduced MDA-MB-231 cells. **B.** Western blots showing NRF2, HIF1α, G3BP1, CAIX and YB-1 levels (relative to GRB2) of MDA-MB-231 cells transduced with *shYB-1 or shScr* vectors grown in ultra-low attachment plates for 24 hours. **C.** Western blots showing HIF1α and YB-1 levels (relative to GAPDH) in *shYB-1* and *shScr*-transduced MDA-MB-231 cells maintained *in vitro* for up to 24 hours in 1% O_2_. **D-E.** Representative images of CAIX (**D**)-, G3BP (**E**)-stained sections of tumours produced from *shYB-1*- or *shScr*-transduced MDA-MB-231 cells. Scale bar, 100 μm. Bar graph shows measured levels of CAIX or G3BP (staining intensity). **F.** Dot plots of tumor sizes obtained in mice injected SQ with T47D cells transduced with GFP-control and *KRAS^G12D^* and assessed 30 days post-transplant. **G.** Representative views of YB-1 levels in the IHC-stained *in vivo* progeny of GFP-control and *KRAS^G12D^*-transduced T47D cells. Bar graph shows quantification of YB-1 expression.

To investigate the possibility that these effects might also be operative during MDA-MB-231-induced tumour formation *in vivo*, we also performed RNAseq, proteomic and immunohistochemical (IHC) analyses of paired transplants of MDA-MB-231 cells transduced with *shYB-1* or control vectors (Appendix Fig S4A-D). This revealed several genes whose encoded RNAs and proteins increased or decreased in concert with reduced levels of YB-1 in transplants generated from cells transduced with the *shYB-1* vector (Appendix Fig S4B-D). These included several genes related to hypoxia (e.g., *PPIF*, *SLC3A2* and *SLC7A1*) (21–23) that are also elevated in breast cancers with increased expression or amplification of *YB-1* (Appendix Fig S4E). Immunostaining of cells generated *in vivo* from the *shYB-1*-transduced MDA-MB-231 cells also showed CAIX and G3BP levels to be decreased compared to the tumours generated from the paired control cells (Fig 3D-E).

To further examine potential effects of deregulated KRAS activity on *YB-1* expression and its associated activities in transformed human mammary cells, we transduced cells from the human T47D breast cancer cell with our lentiviral *KRAS^G12D^* vector (10). As expected, the transduced T47D cells showed the predicted activation of the RAS/RAF/MEK pathway (24) (Appendix Fig S5A) and, in transplanted mice, produced tumours that were larger and grew faster (Fig 3F). IHC analysis of the tumours derived from the transduced cells showed they expressed higher levels of YB-1, G3BP1, HIF1α, and CAIX than control T47D tumours (Fig 3G and Appendix Fig S5B). In addition, when cultured in non-adherent suspension cultures, the *KRAS^G12D^*-transduced T47D cells showed an increased expression of HIF1α and YB-1 compared to controls (Appendix Fig S5C), that was not seen when the cells were cultured *in vitro* under standard adherent 2D conditions (Appendix Fig S5A).

Together, these results suggest that an elevated expression of KRAS in established human breast cancer cells can lead to an enhanced expression of YB-1 and its consequent ability to rapidly activate the expression of stress-ameliorating genes involved in response to hypoxia, and other stresses, and that are required for tumour formation *in vivo*.

### Expression of YB-1 is rapidly increased in tumours produced by normal human mammary cells transduced with *KRAS^G12D^*

We then asked whether elevated *YB-1* expression might be similarly involved in the initial stages of human mammary tumorigenesis driven by experimentally deregulated KRAS activity in freshly isolated normal human mammary cells, as previously described (10). Initial analyses of *KRAS^G12D^*-transduced human mammary cells maintained either in standard 2D adherent cultures for 4 days (Appendix Fig S6A) or in 3D-Matrigel cultures for up to 15 days (Appendix Fig S6B) showed no evidence of altered YB-1 expression. However, 2 weeks after the transduced cells were injected SQ into mice (10^3^ to 2×10^4^ transduced cells in 100 μL of 50% Matrigel/ site), IHC-stained sections of the nascent tumours showed strong increases in YB-1 expression in the cells in comparison to simultaneously stained samples of the matching normal breast tissue samples from which the transduced mammary cells had originally been isolated (Fig 4A and Appendix Fig S6C). In fact, these rapidly increased levels of YB-1 were already similar to those apparent in the larger tumours derived from the same cells but harvested 4-6 weeks later and also stained and analyzed in parallel. This increased YB-1 expression in the tumour cells was confirmed by Western blot analyses of the same samples (Fig 4B).

**Figure 4.**
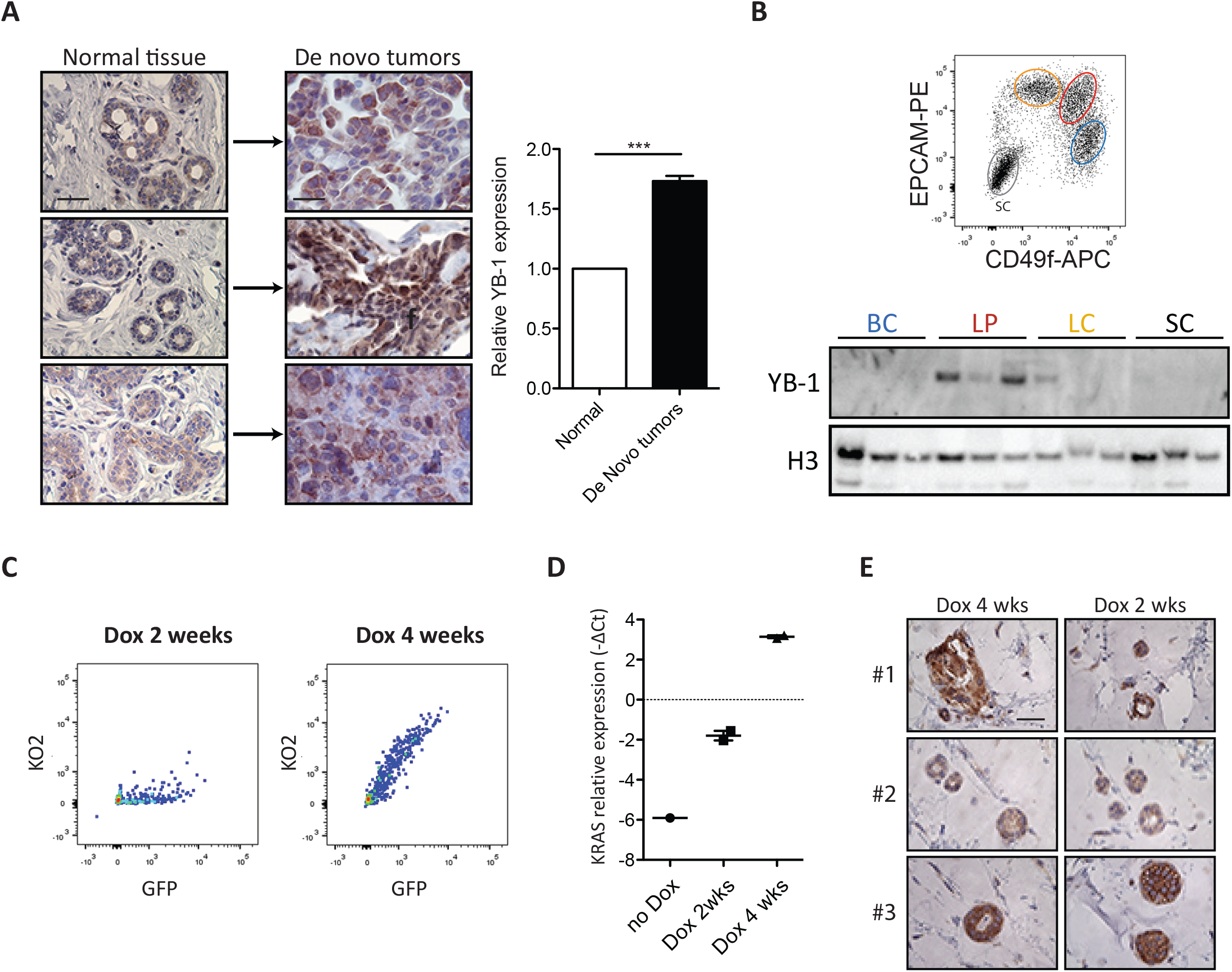
*KRAS^G12D^* upregulates YB-1 in *de novo KRAS^G12D^*-transformed normal human mammary cells. **A** Representative views of YB-1 immunostaining of normal human mammary tissue (left) and 8-week tumours derived from *KRAS^G12D^*-transduced normal mammary cells isolated from the same matching 3 different donors (right). Scale bar, 50 μm. **B** Western blots showing YB-1 levels (relative to H3) in human BCs, LPs, LCs and SCs isolated from 3 normal donors. Subsets were sorted according to their surface EPCAM and CD49f levels (top panel). **C** Representative FACS profile of a 4 week xenograft of inducible *KRAS^G12D^*-transduced human BCs obtained from mice maintained on doxycyline-supplemented water (Dox) for 2 or 4 weeks post-transplant. **D***KRAS* mRNA levels measured in 4 week xenografts of inducible *KRAS^G12D^*-transduced cells obtained from mice maintained on Dox for 0, 2, or 4 weeks, as shown. **E** Representative views of YB-1 immunostaining of 4 week tumours derived from inducible *KRAS^G12D^*-transduced cells in mice maintained on Dox for 2 or 4 weeks (N=3 donors). Scale bar, 50 μm.

We next asked whether sustained expression of KRAS^G12D^ is necessary to maintain the high levels of YB-1 observed in the initial tumour cells appearing *in vivo* from the *KRAS^G12D^*-transduced primary human mammary cells. To address this question, we used a doxycycline (Dox)-inducible *KRAS^G12D^*-Kusabira Orange (KO)-encoding vector (see Methods and Appendix Fig. S6D-E) to transduce freshly purified human mammary cells, and then transplanted these cells SQ as before (4-30×10^4^ cells/site). The injected mice then received Dox either for just the first 2 weeks of a 4-week experiment, or for the entire 4 weeks. At the end of the 4-week period, transplants isolated from the first group of mice contained very few KO+ (*KRAS^G12D^*-expressing) cells (Fig 4C; right panel) and an accompanying large (30-fold) reduction in *KRAS* transcripts 2 weeks later (Fig 4D), as compared to the tumours obtained from mice given Dox for the full 4-week period However, despite the loss of *KRAS* transcripts obtained 2 weeks after discontinuation of Dox, these cells still demonstrated elevated *YB-1* protein levels similar to those evident in the tumours obtained from cells in which *KRAS^G12D^* expression was continuously sustained (Fig 4E). These results indicate that *in vivo*, *KRAS^G12D^* leads to rapidly increased *YB-1* expression in primary cells that can persist even in the absence of continued *KRAS^G12D^* expression.

### Increased AKT-driven *de novo* DCIS is also accompanied by an increase in YB-1 expression, but, on its own increased YB-1 does not confer confer tumorigenic activity

We next designed experiments to determine whether increased expression of YB-1 might be important at the early stage of development of human breast cancers driven by other oncogenes. For this, we transduced separate pools of BCs and LPs from 3 different normal breast tissue samples with mCherry vectors containing a test cDNA, either alone or in combination with a lenti-*KRAS^G12D^*–YFP vector (to look for potential enhancing effects) and a lenti-Luc-YFP vector for bioluminescent monitoring. Cells transduced with the lenti-*KRAS^G12D^*-YFP vector only were included as a positive control and cells transduced with a lenti-Luc-YFP vector only were included as a negative control (Fig 5A).

**Fig. 5.**
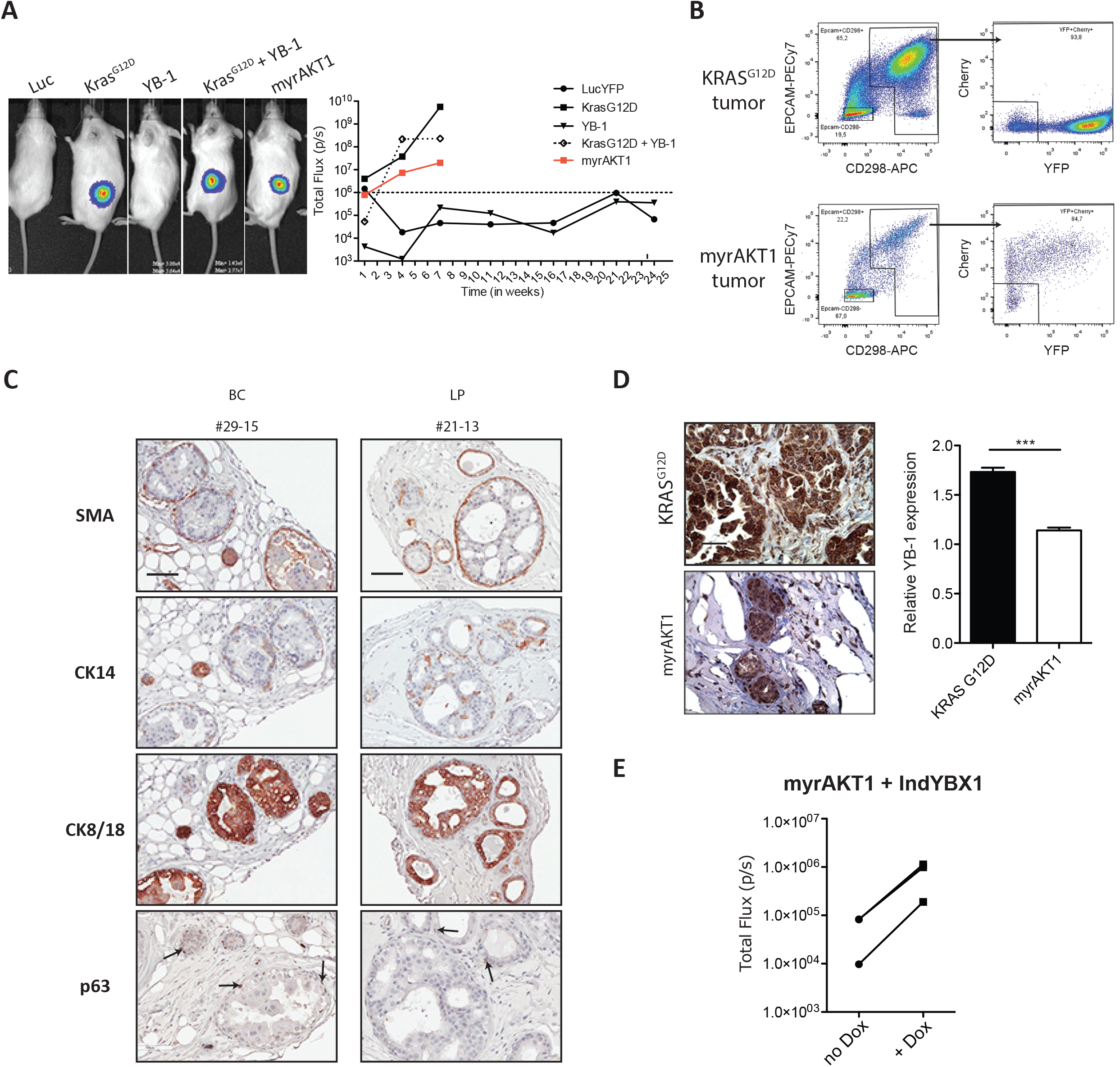
*De novo* formation of DCIS-like tumours leads to modestly increased levels of YB-1. **A** Representative photos of bioluminescence signals measured in mice injected SQ 7 weeks earlier with human mammary cells transduced with lenti-viral vectors encoding Luc-YFP alone or in combination with *KRAS^G12D^*-, YB-1-, *KRAS^G12D^*+YB-1-, or *myrAKT1*. BCs and LPs from 3 donors were pooled separately before transduction. Graph plot shows changes in bioluminescence activity over time. **B** Representative FACS plots of human (CD298/EPCAM)^+^ and mCherry (myrAKT1)^+^ or YFP (KRAS^G12D^)^+^ cells present in dissociated tumours generated from human mammary cells transduced with *KRAS^G12D^* or *myrAKT1*. **C** Representative images of SMA-, CK14-, CK8-18- and p63-stained sections of *myrAKT1*-derived tumours initiated from either BCs or LPs. Scale bar, 100 μm. **D** Representative views of YB-1 immunostaining of 18-week primary *myrAKT1*-derived tumours generated from normal mammary cells from 3 different donors (#1-3). Bar graph shows a comparison of YB-1 staining intensity in *KRAS^G12D^*- or *myrAKT1*-derived tumours. N = 10 (*KRAS^G12D^*) or 6 (*myrAKT1*) tumours. **E** Dot plot of the bioluminescence measured in mice injected SQ with *myrAKT1*+inducible YB-1-transduced human mammary cells and given water with or without doxycyline. N = 3 donors.

Bioluminescence tracking of the transplanted mice showed that ectopic expression of YB-1 alone was insufficient to confer tumour-forming capacity on primary human mammary cells and, when combined with *KRAS^G12D^* in the same cells, also did not enhance the growth of the tumours generated from cells transduced with *KRAS^G12D^* only (Fig 5A). In these experiments, we also confirmed that on their own, neither dominant-negative forms of *TP53*, nor a mutant form of *PI3K* were able to induce the cells to become tumorigenic (10). Negative results were also obtained for vectors encoding *EGFR* and *c-MYC*, and shRNAs targeting *PTEN* and *BRCA1* transcripts (data not shown). However, forced expression of a cDNA encoding *myrAKT* alone, led to significantly increasing luciferase signals over an 8-week period post-transplant, albeit at consistently lower levels than those obtained from *KRAS^G12D^*-transduced cells (Fig 5A). Constitutive activity of the myrAKT protein in this model system is attributed to loss of the pleckstrin homology domain and addition of an engineered SRC myristoylation signal sequence that targets the protein to the cell membrane (25). Phenotypic analysis of tumours obtained from recipients of *myrAKT*- transduced cells with showed that they were universally human EpCAM^+^CD298^+^ as well as mCherry^+^, indicative of an oncogenic role of deregulated AKT1 activity in these cells (Fig 5B). The modest tumorigenic activity of *myrAKT* was also readily replicated (9/13 tests) in either transduced BCs or LPs obtained from different donors (Appendix Fig S7A).

Histological analysis of the tumours produced from the *myrAKT1*-transduced cells showed that they closely resemble DCIS. This included a confined organization of the cells in duct-like structures with an outer layer of cells with basal features (expression of smooth muscle actin and TP63) (Fig 5C and Appendix Fig S7B), and extensive luminal filling by cells with luminal features (strong CK14 and CK8/18 positivity) (Fig 5C and Appendix Fig S7B). Tumours also contained a low frequency of ER^+^ and/or Ki67^+^ cells, and PR^+^ cells were not detected (Appendix Fig S7C). Importantly, these structures also showed increased YB-1 protein as compared to normal cells, but less than the YB-1 levels typical of the cells in the *KRAS^G12D^*-induced tumours (Fig 5D). Interestingly, mice transplanted with cells co-transduced with the Dox-inducible KO/YB-1 vector (Appendix Fig S8A) plus the lenti-*myrAKT*-mCherry vector, when treated with Dox, showed increased luciferase activity (Fig 5E), along with increased YB-1 expression, compared to matched transplanted injected into mice that were not treated with Dox (Appendix Fig S8B).

These experiments extend the range and genetic determinants of *de novo* transformed primary human mammary cell types that show upregulation of YB-1 to now include a DCIS model that does not involve deregulation of KRAS. At the same time, they show that a forced increase in YB-1 expression can enhance the growth of DCIS tumours in the *myrAKT* model, although again, increased YB-1 alone did not confer tumorigenic activity in this setting.

### *KRAS^G12D^* requires YB-1 to induce its tumorigenic activity and the HIF1α response in transduced normal human mammary cells

IHC analysis of *de novo* tumours derived from *KRAS^G12D^*-transduced human mammary cells showed that they contained increased levels of stress-related translational targets of YB-1, including HIF1α, NRF2, G3BP1 and membranous CAIX (Fig 6A), compared to matching normal tissue. To determine whether YB-1 is *required* for the *in vivo* genesis of tumours from *de novo* transformed human mammary cells, we used the same knockdown strategy as with the MDA-MB-231 cells. Accordingly, purified BCs and LPs were transduced first with *shYB-1* or control (*shScr*) lentiviral vectors (Appendix Fig S9A) and then, after a further 2 days *in vitro*, the cells were transduced at high efficiency with vectors encoding both *KRAS^G12D^*-mCherry and a luciferase (Luc)-YFP vector prior to being transplanted SQ into immunodeficient mice (2×10^4^-2.5×10^5^ cells/transplant). Two weeks following transplantation, cells derived from either BCs or LPs that had been co-transduced with *KRAS^G12D^* and *shYB-1* produced 2-1,000-fold lower bioluminescent signals than those obtained from their counterparts transduced with *KRAS^G12D^* and the control sh vector (Fig 6B). H&E-staining showed that the cellularity of the BC and LP-derived tumors was greatly reduced compared to that of parallel tumours generated from the control cells (cells transduced with *KRAS^G12D^* plus the *shScr* construct, Appendix Fig S9B). Because of the small size of the *KRAS^G12D^* + *shYB-1*-derived tumors, we limited our studies on potential YB-1 cytoprotective effects on just the HIF1α pathway in these tumours. IHC staining confirmed that the total number of YB-1^+^ cells was reduced in the test cell-derived harvests (Fig. 6C) as were levels of HIF1α (Fig 6D), CAIX (Fig 6E), VEGF (Fig 6F) and CD34 (the latter as a marker of blood vessels; Appendix Fig S9C).

**Figure 6.**
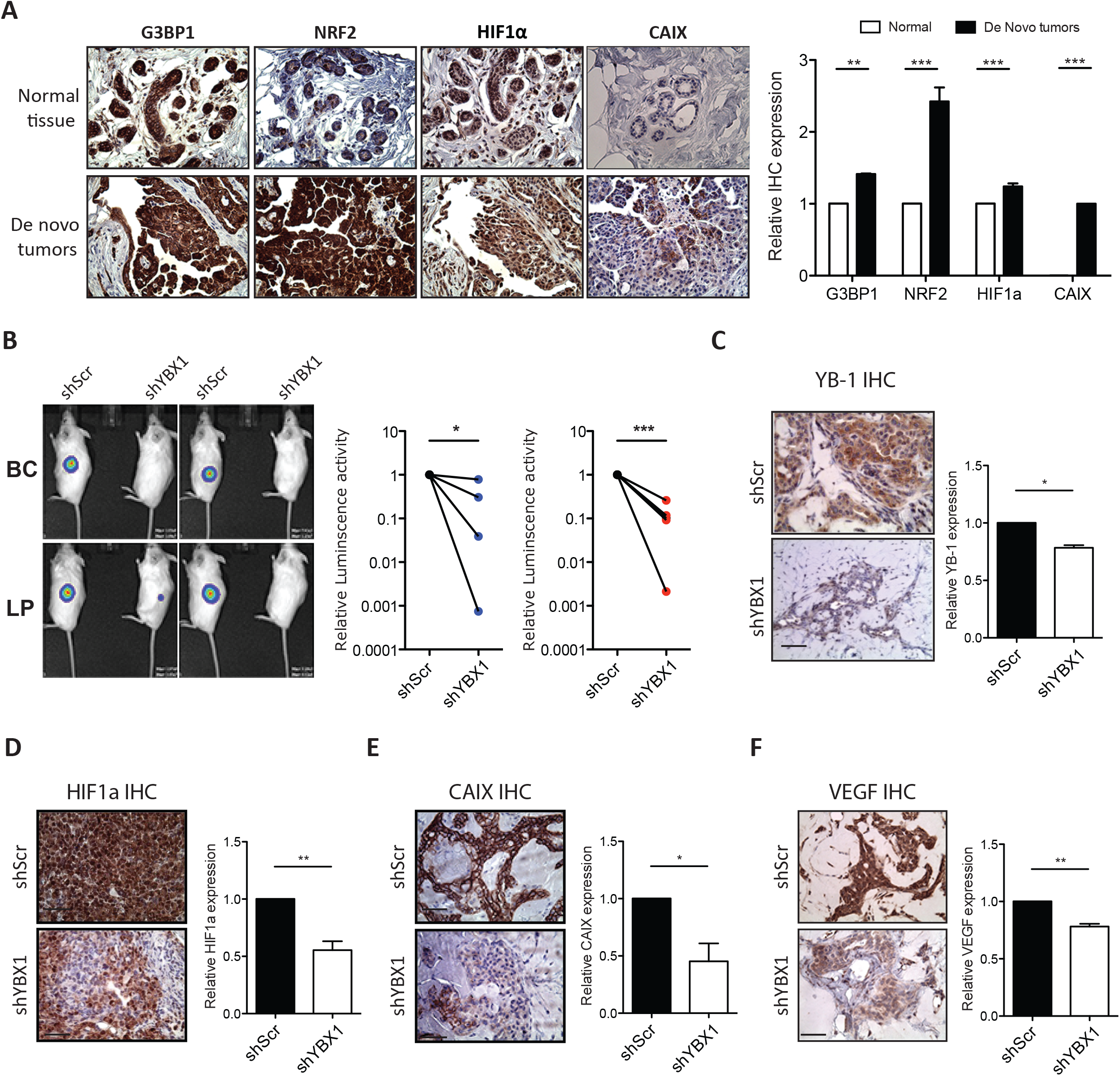
Increased YB-1 is necessary for *KRAS^G12D^*-induced human mammary tumorigenesis. **A** Representative views of G3BP1, NRF2, CAIX and HIF1α immunostaining of normal human mammary tissue (left) and 8-week tumours derived from *KRAS^G12D^*-transduced normal mammary cells isolated from the same matching 3 different donors (right). Scale bar, 50 μm. Bar graph shows quantification of G3BP1, NRF2, HIF1α and CAIX expression. **B** Representative photos of the bioluminescence signals obtained in mice injected SQ with *KRAS^G12D^*+*shYB-1*- or *KRAS^G12D^*+*shScr*-transduced primary human mammary cells 2 weeks earlier. Dot plot showing the bioluminescence activity of tumours derived from BCs (blue) and LPs (red). **C-F** Representative images of YB-1 (**C**)-, HIF1α (**D**)-, CAIX (**E**)- and VEGF (**F**)-stained sections from different BC- or LP-derived tumours arising from *KRAS^G12D^*+*shYB-1*- or *KRAS^G12D^*+*shScr*-transduced cells. Scale bar, 100 μm. Bar graphs shows quantification of YB-1 (**C**)-, HIF1α (**D**)-, CAIX (**E**)- and VEGF (**F**)-intensities in tumours derived from *KRAS^G12D^*+*shYB-1*- or *KRAS^G12D^*+*shScr* -transduced cells. N = 8. P-values in panels (A-E) were determined by Student’s t-test, \**P*<0.05, \*\**P*<0.01, \*\*\**P*<0.001.

Together, these experiments establish that the initial tumorigenic activity displayed by xenografts of normal human mammary cells forced to express *KRAS^G12D^* is highly dependent on an upregulation of YB-1 and this is associated with a YB-1-dependent expression of proteins that mediate stress responses.

## DISCUSSION

Here we show that increased expression of *YB-1* characteristic of patients’ breast cancers with poor prognosis is associated with elevated KRAS or AKT activity and an activated stress response program. In addition, we show that these associations drawn from patient data are reproduced in several experimental models of human breast cancer growing *in vivo* in transplanted immunodeficient mice. The latter include the established MDA-MB-231 human breast cancer cell line that has a *KRAS^G13D^* mutation (18) and highly penetrant metastatic as well as tumorigenic activity in transplanted immunodeficient mouse hosts. In addition, we show that upregulation of YB-1 protein is a rapidly acquired feature of tumours produced in mice transplanted with T47D cells or primary normal human mammary cells forced to overexpress *KRAS^G12D^*. We also describe a new, less aggressive model of *de novo* normal human mammary cell transformation with features of DCIS that results from lenti-virally deregulated AKT activity, and also displays increased expression of YB-1. Taken together, these findings indicate a role for YB-1 both at the point of initial transformation of normal human mammary cells, as well as in more advanced derivatives that display metastatic activity in transplanted immunodeficient mice.

Previous experiments in which YB-1 was overexpressed in the immortalized but non-tumorigenic MCF10A cell line transformed with HRAS showed that high YB-1 expression contributes to the disruption of the normal mammary cell architecture they produce and promotes an epithelial mesenchymal transition (9). However, published patient breast cancer datasets (14, 15, 26), show a clear link between increased *YB-1* expression and amplified *KRAS*, but no association with *HRAS* or *NRAS*. Here we provide both *in vitro* and *in vivo* evidence of a similar loss of a normal acinar architecture by *KRAS^G12D^*-transduced human mammary cells, and a requisite increase in their expression of *YB-1* and its associated activated stress response targets. These include results obtained from 3D cultures of both T47D cells and primary human mammary cells (not seen in standard 2D conditions), and in tumours produced in mice transplanted with these cells. Our findings thus add new weight to a growing body of evidence that induced expression of YB-1 confers a cytoprotective mechanism to transformed cells exposed to increased stresses, such as those created during tumour growth or metastasis, or growth under certain adverse conditions *in vitro*.

This concept is further supported by the specific YB-1-associated increases in G3BP1, NRF2, HIF1α and CAIX evident in the tumourigenic systems studied here. Notably, our results showing that suppression of YB-1 leads to decreased expression of HIF1α and CAIX in both *de novo* and advanced tumours, extend the finding of heightened expression of HIF1α and HIF target genes in patients’ breast cancers with amplified KRAS. Together, these findings support a model in which activated KRAS or AKT leads directly or indirectly to increased levels of YB-1 levels which, in turn, promote an elevated HIF1α response and potentially other cytoprotective programs required for tumour *in vivo*. Importantly, the shared high expression of *HIF1α* and its target genes in triple-negative breast cancers (27) underscores the importance of our findings in several model systems and the opportunities they afford for future mechanistic and translational investigations. Notably, we have previously shown that HIF1α induction in sarcoma cells is mediated by direct binding of YB-1 to the 5’-UTR of *HIF1A* transcripts to directly enhance their translation (5). In addition, we have also demonstrated that YB-1 regulates stress granule formation and sarcoma tumor progression by translationally activating G3BP1 and SG formation (8). KRAS has also been shown to promote SG formation following exposure colon and pancreatic cancer cells to stress-inducing stimuli (7). YB-1 might thus represent a common intermediate in the activation of many cytoprotective programs that are required for multiple types of tumour cells to survive and grow when they are exposed to different stress conditions *in vivo*.

Our findings reported here using *de novo* models of human DCIS and IDC suggest that YB-1 plays a critical role in initiating the growth of transformed human mammary cells *in vivo* with accompanying activation of downstream targets involved in stress response programs, thus extending findings associated with the growth of more aggressive and metastatic breast cancer cells. YB-1 may therefore represent a potential relevant target for therapeutic intervention in breast cancer. We recently reported that class I histone deacetylase (HDAC) inhibitors induce hyperacetylation within the RNA binding CSD of YB-1 in sarcoma cells (6), thus reducing its target mRNA binding activity and downstream effects on their translational activation. At the same time, the present findings suggest that targeting HIF1α or its downstream effectors may represent alternative and potentially more tractable clinical targets. The present studies indicate how these issues might now be investigated in a range of relevant human breast cancer models that include *de novo* transformants of primary cells created with defined oncogenic perturbations.

## MATERIALS AND METHODS

### Cells and cultures

Normal human reduction mammoplasty discard tissue was collected with informed consent, according to protocols approved by the University of British Columbia Research Ethics Board. Organoid-rich pellets were then isolated and viably cryopreserved (28). As required, thawed organoids were rinsed with 2% fetal bovine serum (FBS, from STEMCELL Technologies) in Hank’s Balanced Salt Solution (HF), and the cells then dissociated in 2.5 mg/ml trypsin with 1 mM EDTA and 5 mg/ml dispase (STEMCELL Technologies) with 100 μg/ml DNaseI (Sigma) and washing of the cells with HF between each step. The resulting cell suspension was filtered through a 40 μm mesh and BCs then isolated by FACS according to their CD45^−^CD31^−^EpCAM^lo^CD49f^+^ phenotype, LPs according to their CD45^−^CD31^−^EpCAM^hi^CD49f^+^ phenotype, LCs according to their CD45^−^CD31^−^EpCAM^hi^CD49f^−^ phenotype and stromal cells (SCs) according to their CD45^−^CD31^−^EpCAM^−^CD49f^−^ phenotype using well established protocols and reagents (29). Following FACS, cells were transduced or cultured in SF7 media supplemented with 5% FBS. MCF10A cells (obtained from J Brugge, Harvard University, Cambridge, MA) were maintained in phenol-free DMEM/F12 nutrient mix supplemented with 5% horse serum, 10 mg/ml insulin, 0.5 mg/ml hydrocortisone, 100 ng/ml cholera toxin, 20 ng/ml EGF (all Sigma), and 1% penicillin/streptomycin (Life Technologies). 3D assays of human mammary cells were performed by culturing the cells in the presence of irradiated 3T3 fibroblasts for 8, 10 or 14 days in Matrigel (Corning) SF7 media supplemented with 5% FBS as previously described (30). MDA-MB-231 cells were obtained from S. Dunn (Child and Family Research Institute, Vancouver, BC) and maintained in DMEM with 10% FBS. Their identity was confirmed by DNA sequencing, including detection of the *KRAS^G13D^* allele (18). T47D cells were obtained from Joanne Emerman (Department of Anatomy, University of British Columbia) and maintained in RPMI with 10% FBS. For 3D cultures, cells were transferred to ultra-low attachment surface plates and cultured for 24 hours before harvesting.

### Transduction and transfection

Primary cells were transduced with lenti-viral vectors prepared and used as previously described(10). For stable inhibition of YBX1 transcripts, lentiviral vectors encoding sh*YBX1* (sc-38634-V, Santa Cruz, #1307/TRCN0000315307, #1309/TRCN0000315309, and #948/TRCN0000007948, Sigma), or sh*Scr* (sc-108080, Santa Cruz or shScr (SHC202, Sigma) were purchased or constructed.

### Xenografts

Female immunodeficient NOD-*Rag1^−/−^IL2Rγc^−/−^* (NRG) mice were bred and housed in the animal facility at the British Columbia Cancer Research Centre under SPF conditions. Surgery was performed on 5- to 10-week-old mice. All experimental procedures were approved by the University of British Columbia Animal Care Committee. Mice that have been xenografted with cells containing inducible vectors have been supplied with Dox (1mg/ml) dissolved in water.

To generate primary tumours, enzymatically dissociated human mammary cell suspensions were prepared, transduced and transplanted into mice SQ in 50% (v/v) Matrigel (10). To measure tumour bioluminescence from the co-transduced luciferase cDNA, mice were injected intraperitoneally with 150 mg/kg body weight of d-luciferin (Promega) and 10 minutes later the mice were imaged using a Xenogen IVIS Lumina system with Living Image version 3.0 software (Caliper Life Sciences). To prepare single cell suspensions from excised tumours, the tissue was minced with a scalpel, incubated at 37 °C in DMEM/F12 media supplemented with 5% FBS and 300 U/ml collagenase and 100 U/ml hyaluronidase for 1 to 2 hours with periodic vortexing, washed with HF, and treated with 2.5 mg/ml trypsin with 1 mM EDTA and 5 mg/ml dispase with 100 μg/ml DNaseI. Human cells were sorted after staining with anti-human specific antibodies directed against EpCAM and CD298 (Biolegend) with simultaneous depletion of mouse cells stained with anti-mouse-specific antibodies directed against CD45 and CD31 (Biolegend).

### IHC staining

Pieces of tumours obtained from mice or normal breast were fixed in 10% buffered formalin (Fisher), washed in 70% ethanol and embedded in paraffin. Sections of paraffin-embedded tissue (3 mm) were first treated with Target Retrieval solution (DAKO) and then a cytomation serum-free protein block (DAKO) followed by staining with specific antibodies recognizing human YB-1 (#HPA040304, Sigma), ER (SP1; 1/50; Thermofisher; RM9101), PR (SP2; 1/50; Neomarker; 9102), Ki67 (SP6; 1/50; Thermofisher; RM9106), CK14 (Novocastra/Leica; 1/50; NCL-L-LL02), CK8/18 (Novocastra/Leica; 1/50; NCL-L-5D3), p63 (4A4; 1/50; Gentex; GTX23239), SMA (1A4; 1/100; Dako; MO851), HIF1a (Novus Biologicals; 1/150; NB100-134), CAIX (R&D Systems; 1/200; AF2188), CD34 (Abcam; 1/400; Ab81289), NRF2 (Abcam. 1/150; Ab62352), G3BP (Novus Biologicals; 1/500; NBP2-16563). A secondary mouse or rabbit antibody conjugated to horseradish peroxidase and treatment with 3,3ʹ-diaminobenzidine (DAB, DAKO) was used to obtain a positive brown staining. Negative IgG controls were performed on normal reduction mammoplasty tissue.

Quantitative analysis of IHC samples was conducted using the colour deconvolution plugin which implements stain separation and the ImmunoRatio plugin for ImageJ software (developed at the National Institutes of Health, USA, and available at http://rsb.info.nih.gov/ij/). Student’s t-test was used for data analysis, unless indicated otherwise.

### Plasmids

Inducible *KRAS^G12D^*- and YB-1-encoding vectors were generated in a pINDUCER21 backbone (31) by replacing the attR1-ORF-attR2 cassette with a KRAS-2A-KO2 fragment.

### WB and densitometry analysis

After the required treatment, cells were washed with cold PBS and incubated for 15 minutes at 4°C with RIPA lysis buffer (30 mM Tris-HCl, pH 7.5, 150 mM NaCl, 10% glycerol, 1% Triton X-100 (Sigma) supplemented with a 1 mM NaF, 1 mM NaVO3 and 1 mM PMSF (all Sigma). Cells extracts were centrifuged at 13,000 g for 10 minutes at 4°C. The protein concentration of the supernatant fraction was determined using the Bio-Rad Bradford Protein Assay Kit according to the manufacturer’s instructions. For each sample, an equal amount of total protein was diluted in sample buffer (Invitrogen) and boiled for 5 minutes. Samples were loaded onto precast NuPAGE 4-12% polyacrylamide gels (Invitrogen). After electrophoresis, proteins were transferred to a PVDF transfer membrane. Membranes were then blotted overnight at 4°C with appropriate primary antibodies, such as anti-ACTIN (Santa Cruz, sc-1615, 1/10,000), anti-H3 (Cell Signaling Technology, 12648, 1/10,000), anti-RAS (Cell Signaling Technologies, 3339, 1/1,000), anti-YB-1 (Cell Signaling Technology, 4202, 1/1,000) and anti-HIF1a (Cayman; 10006421; 1/1000). Specific binding of antibodies was detected using appropriate secondary antibodies conjugated to horseradish peroxidase, and visualized with SuperSignal™ West Femto Maximum Sensitivity Substrate (Thermofisher) on a ChemiDoc Gel Imaging system (Bio-rad). Densitometric analyses of immunoblots were performed using ImageJ.

### RNAseq data

Copy number alterations and Z-score normalized RNAseq expression values (V2 RSEM) were obtained from cBioPortal (32), from TCGA (13), METABRIC (14) and Mutational profiles of metastatic breast cancers (15) datasets. Paired-end reads were generated from MDA-MB-231 shScr or shYBX1 cells or derived tumors, on an Illumina HiSeq2500 sequencer. Read sequences were aligned to the hg19 human reference using the BWA-SW algorithm (33) to generate binary alignment/map (BAM) files. Transcript counts were obtained with the summarize Overlaps function from GenomicAlignments package(34). Differential expression analysis was performed with DESeq2 package (35).

### Proteomic data

Tissues were thawed and lysed in 100 μL lysis buffer containing 500 mM Tris-HCL pH 8, 2% SDS (w/v), 1% NP-40 (v/v), 1% Triton X100 (v/v), 0.5 mM EDTA, 50 mM NaCl, 10 mM Tri(2-carboxyethyl)phosphine (TCEP) and 40 mM chloroacetamide (CAA). The proteins were then denatured by heating at 95°C for 90 minutes with shaking at 1,100 rpm before incubation at room temperature for 90 minutes in the dark to allow reduction and alkylation of disulfide bonds by TCEP and CAA respectively. SP3 beads (36, 37) were added and the tissues were sonicated in a Bioruptor Pico (Diagenode) for 10 cycles (30 seconds ON, 30 seconds OFF). The samples were purified and prepared for trypsin digestion using the SP3 method (37). Tryptic peptides from each sample were individually labeled with TMT 10-plex labels (Thermo Scientific), pooled, and fractionated into 12 fractions by high pH RP-HPLC, desalted, and then analyzed using an Easy-nLC1000 liquid chromatograph (LC) (Thermo Scientific) coupled to a Orbitrap Fusion Tribrid mass spectrometry (MS) (Thermo Scientific) operating in MS3 mode. The offline peptide fractionation and LC-MS conditions are as described (37). The raw MS data were searched using Proteome Discoverer (version 2.1.1.21) using the embedded Sequest HT algorithm against a combined UniProt Human proteome database with a list of common contaminants appended (24,624 total sequences). Sequest HT parameters were specified as: trypsin enzyme, allowance for 2 missed cleavages, minimum peptide length of 6, precursor mass tolerance of 20 ppm, and a fragment mass tolerance of 0.6. Dynamic modifications allowed were oxidation of methionine residues, and TMT at lysine residues and peptide N-termini. Carbamidomethylation of cysteine residues was set as a static modification. Peptide spectral match (PSM) error rates were determined using the target-decoy strategy coupled to Percolator modeling of positive and false matches (38, 39). Data were filtered at the PSM-level to control for false discoveries using a q-value cutoff of 0.05 as determined by Percolator. Contaminant and decoy proteins were removed from all datasets prior to downstream analysis. Statistical analysis of differential protein expression was performed at the peptide level using a modified version of the PECA function that is appropriate for input of log-transformed data (40). PECA uses Limma (41) to generate a linear model for estimating fold changes and standard errors prior to empirical Bayes smoothing. Median t-statistics of the assigned peptides were used to calculate false-discovery rate-adjusted *p*-values determined from the beta distribution, as described previously (40).

### RT-PCR

Total RNA was extracted from cryopreserved tumour samples or cultured cells using the Total RNA Isolation Micro kit (Agilent) and cDNA then synthesized using SuperScript VILO cDNA synthesis kit (Life Technologies). RT-PCR was performed using a SYBR Green master mix (Applied Biosystems) and samples run in triplicate with custom-designed primers.

### Statistical analyses

Values are expressed as mean ± SEM, unless otherwise specified. Significance was evaluated using Student’s t-test, unless otherwise specified. \**P*<0.05, \*\**P*<0.01, \*\*\**P*<0.001 ns = not significant.

## Supporting information

Supplemental informations

## ACKNOWLEDGMENTS

The authors thank D. Wilkinson, G. Edin, M. Hale, and A. Li for excellent technical support; and to Drs. E. Bovill, J. Boyle, S. Bristol, P. Gdalevitch, A. Seal, J. Sproul, and N. van Laeken for access to discarded reduction human mammoplasty tissue. This work was supported by grants to CJE from the Canadian Cancer Society Research Institute, the Cancer Research Society, and the Cancer Institute for Health Research (CIHR grant CRP-154482), to PHS from the Terry Fox Research Institute (Team Grant 1021) and the CIHR (Foundation 143280). ST held a CIHR Banting and Best Studentship and A.E-N a Michael Smith Foundation for Health Research trainee award (# 17159).

## AUTHOR CONTRIBUTIONS

S.L., P.H.S. and C.J.E. conceptualized this project and drafted the manuscript. S.L., A.E-N., S.T., and S.C. performed experiments. S.C and G.L.N. carried out the computational and bioinformatics analysis of the proteomics and RNA-seq data. S.L., A.E-N., S.T., S.C., G.L.N., M.H., B.G., G.B.M., P.H.S. and C.J.E. analyzed and interpreted the data.

