## Supplemental informations for "YB-1 IS REQUIRED FOR THE GENESIS AND METASTATIC CAPACITY OF HUMAN BREAST CANCER"

#### Supplementary Figure Legends

**Appendix Figure S1. High levels of *YBX1* transcripts are associated with metastatic human breast cancers and *KRAS*-amplified tumors.**

**A-B**. *YBX1* mRNA expression according to the ER (**B**) or metastatic status (**C**) of invasive breast carcinomas in the TCGA dataset.

**C.** *YBX1* mRNA expression according to gene copy numbers of *TP53, ERBB2*, *PIK3CA* and *AKT1* in invasive breast carcinomas in the TCGA dataset. All *YBX1* values are shown as RPKMs.

**D.** Representative table of *YBX1* alteration co-occurrence in metastatic breast cancer.

**Appendix Figure S2. *KRAS* and AKT1 amplifications are associated with a YB-1-regulated stress response.**

**A.** *VEGFA* (left panel) and *NFE2L2* (right panel) mRNA expression according to *KRAS* copy number status in invasive breast carcinomas in TCGA dataset. Values for *VEGFA, and NFE2L2* are shown as RPKMs.

**B.** *CAIX* (left panel), and *VEGFA* (right panel) transcript levels (RPKM values) shown according to *KRAS* copy number status in the invasive breast carcinoma data set in METABRIC.

**C.** Scatter plot of *YBX1* and *CAIX* mRNA expression in amplified-*KRAS* invasive breast carcinomas (METABRIC dataset).

**D.** *HIF1A* (left panel), *CAIX* (middle panel), and *VEGFA* (right panel) transcript levels are shown according to *YBX1* copy number status in the TCGA invasive breast carcinoma dataset. All transcript values are shown as RPKMs.

**E.** *VEGFA* (left panel) and *CAIX* (right panel) transcript level (RPKM values) shown according to *AKT1* copy number status in the invasive breast carcinoma data set in METABRIC.

**F.** *YBX1*, *HIF1A*, *CAIX* and *VEGFA* transcript levels are shown according to *AKT1* copy number status in the TCGA invasive breast carcinoma dataset. All transcript values are shown as RPKMs.

**G-H**. Scatter plot of *YBX1* and *CAIX* mRNA expression in amplified-*AKT1* invasive breast carcinomas in METABRIC (**C**) or (**D**) TCGA datasets.

**Appendix Figure S3. YB-1 is required for dissemination of intravenously injected MDA-MB-231 cells.**

**A**. YB-1 expression analyzed by Western blot assessment of extracts of sh*YBX1*- or sh*Scr*-transduced MDA-MB-231 cell harvests.

**B**. YB-1 expression analyzed by FACS analysis of individual sh*YBX1*– or sh*Scr*-transduced MDA-MB-231 cell harvests.

**C**. Representative pictures of bioluminescence signals in mice injected subcutaneously with MDA-MB-231 cells transduced with sh*Scr* or sh*YBX1*. Dot plot shows the measured bioluminescence in these tumours 26 days post-transplant.

**D**. Weights of the tumours shown in (**C**).

**E.** Representative views of YB-1 levels in the IHC-stained *in vivo* progeny of sh*YBX1*- or sh*Scr*-transduced MDA-MB-231 cells. Bar graph shows quantification of YB-1 expression.

**F.** Representative pictures 45 days post-transplant of bioluminescence signals measured in mice injected intravenously with sh*YBX1*- or sh*Scr*-transduced MDA-MB-231 cells. Dot plot showing levels of bioluminescence of tumors derived from MDA-MB-231 cells transduced with sh*Scr* or a sh*YBX1* and assessed 47 days post-transplant.

**G.** Representative H&E-stained photomicrographs of lungs of mice injected intravenously with sh*YBX1*- or sh*Scr*-transduced MDA-MB-231 cells.

**H.** Comparison of YB-1 levels (staining intensity) in lung tumors shown in **G**.

**Appendix Figure S4. Inhibition of *YBX1* expression downregulates expression of *PPIF*, *SLC3A2* and *SLC7A1*.**

**A**. Hierarchical clustering of proteomics data obtained on tumour cells generated from sh*YBX1- or* sh*Scr*-transduced MDA-MB-231 cells.

**B**. Correlation plot between transcripts and proteins identified by RNAseq and proteomic analysis of tumours derived from sh*YBX1*- or sh*Scr*-transduced MDA-MB-231 cells.

**C-D.** Genes whose transcript and protein expression in MDA-MB-231 cells after transduction with sh*YBX1* was either upregulated (**C**) or down-regulated (**D**) by comparison to sh*Scr*-transduced (control) cells from analyses of matched pairs of RNAseq and proteomics data.

**E.** *SLC3A2, PPIF* and *SLC7A1* mRNA expression according to *YBX1* copy number status in invasive breast carcinomas in the TCGA data. Values for *SLC3A2* and *PPIF* are shown as RPKMs.

**Appendix Figure S5**. ***KRAS*$^{G12D}$-overexpression lead to YB-1 increase *in vitro* only in stressed conditions.**

**A** Western blots of GFP-control and *KRAS*$^{G12D}$-transduced T47D cells, showing YB-1, P-ERK1/2, ERK1/2 and RAS (relative to Actin). Dot plots showing P-ERK (relative to Actin) and YB-1 (relative to Actin) levels.

**B** Representative views of G3BP1, HIF1α and CAIX immunostaining in the IHC-stained *in vivo* progeny of GFP-control and *KRAS*$^{G12D}$-transduced T47D cells. Scale bar, 50 μm. Bar graph shows quantification of G3BP1, HIF1α and CAIX expression.

**C** Western blots showing HIF1α, YB-1 and RAS levels (relative to GAPDH) of GFP-control and *KRAS*$^{G12D}$-transduced T47D cells grown in ultra-low attachment plates for 48h.

**Appendix Figure S6**. ***KRAS*$^{G12D}$-transduced primary human mammary cells display high levels of YB-1 only *in vivo*.**

**A.** Western blots showing YB-1 and RAS levels (relative to H3) in control and *KRAS*$^{G12D}$-transduced human BCs and LPs from 3 normal donors, as assessed 3 days post-transduction.

**B** Representative immunofluorescence images of control and *KRAS*$^{G12D}$-transduced BCs (top) and LPs (bottom) assessed 15 days post-transduction, and cultured in 3D in Matrigel. Staining was performed using an anti-YB-1 antibody, Phalloidin (F-ACTIN) and DAPI (DNA).

**C**. Representative images of YB-1 immunostaining of 2 week-old xenografts of

*KRAS*$^{G12D}$-transduced primary human mammary cells. Scale bar, 200 µm (left) or 100 µm (right).

**D.** Representative FACS profile of human mammary MCF10A cells stably expressing an inducible KRAS-2A-KO2 construct after being maintained in the presence or absence of doxycycline.

**E.** Western blots showing RAS levels (relative to ACTIN) in the same cells as in (**D**).

**Appendix Figure S7. Transplants of primary human mammary cells expressing *myrAKT1* produce DCIS-like structures.**

**A**. Representative photos of bioluminescence signals obtained in mice injected subcutaneously 5 or 7 weeks earlier with Luc-YFP- and *myrAKT1*-transduced human mammary cells. Graph plot shows bioluminescence activity from tumours derived from BCs (blue) and LPs (red). N = 5 donors.

**B-C**. Representative IHC images of sections stained for SMA, CK14, CK8-18 and p63 (**B**); or ER, PR, and Ki67 (**C**) in tumours derived from transplants of either BCs or LPs-transduced with *myrAKT1*. Scale bar, 100 µm.

**Appendix Figure S8. YB-1 induction model.**

**A.** Representative FACS profile (left panel) and Western blots (right panel) of human mammary MCF10A cells expressing inducible YBX1-2A-KO, cultured with or without doxycycline.

**B.** Representative IHC images of sections stained for YB-1 in tumors generated from LPs (left) or BCs (right) transduced with *myrAKT1* vector an inducible YBX1, and transplanted into mice given water with (bottom) or without doxycycline (top).

**Appendix Figure S9. Suppression of YB-1 expression impairs *in vivo* tumour formation by *KRAS*$^{G12D}$-transduced normal human mammary cells.**

**A.** Western blot showing YB-1 expression in cells expanded *in vitro* from isolated BCs and LPs transduced with sh*YBX1 or* sh*Scr* using cells from 3 normal donors.

**B.** Representative images of H&E-stained sections from different BC- or LP-derived tumours arising from *KRAS*$^{G12D}$+sh*YBX1*- or *KRAS*$^{G12D}$+sh*Scr*-transduced cells. Scale bar, 200 μm.

**C.** Representative images of CD34-stained sections from different BC- or LP-derived tumours arising from *KRAS*$^{G12D}$+sh*YBX1*- or *KRAS*$^{G12D}$+sh*Scr*-transduced cells. Scale bar, 200 μm. Bar graphs shows quantification of CD34 intensities in tumours derived from *KRAS*$^{G12D}$+sh*YBX1*- or *KRAS*$^{G12D}$+sh*Scr*-transduced cells from 6 normal donors.

### Appendix Figure S1

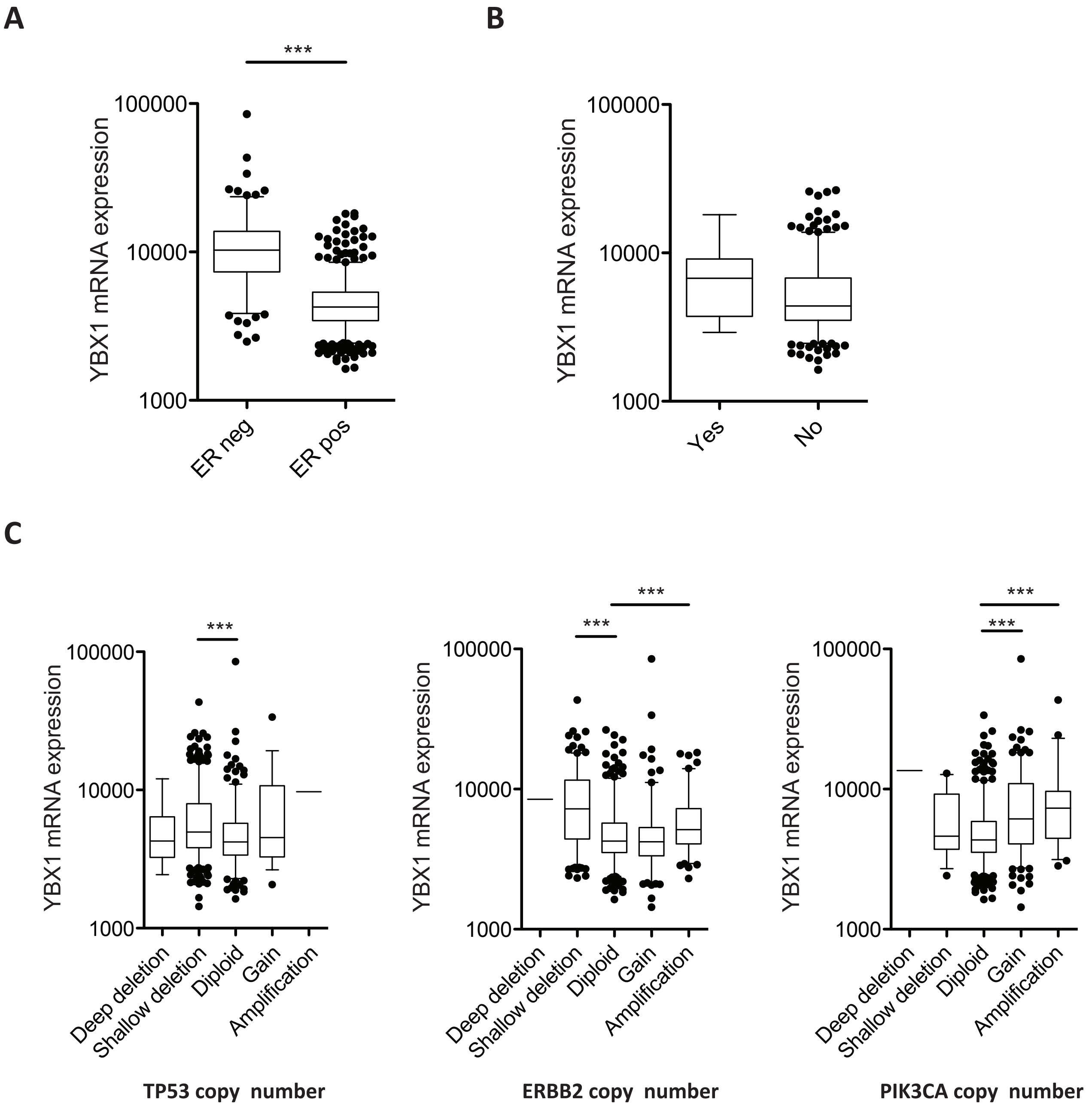


**D**

YBX1 co-occurrent alterations

| Gene | p-value | Log odds ratio |
|---|---|---|
| KRAS | 0,026 | 2.71 |
| TP53 | 0,150 | 1.43 |
| PIK3CA | 0,609 | -0.37 |
| ERBB2 | 0,369 | 1.04 |

**Metastatic Breast Cancer**

### Appendix Figure S2

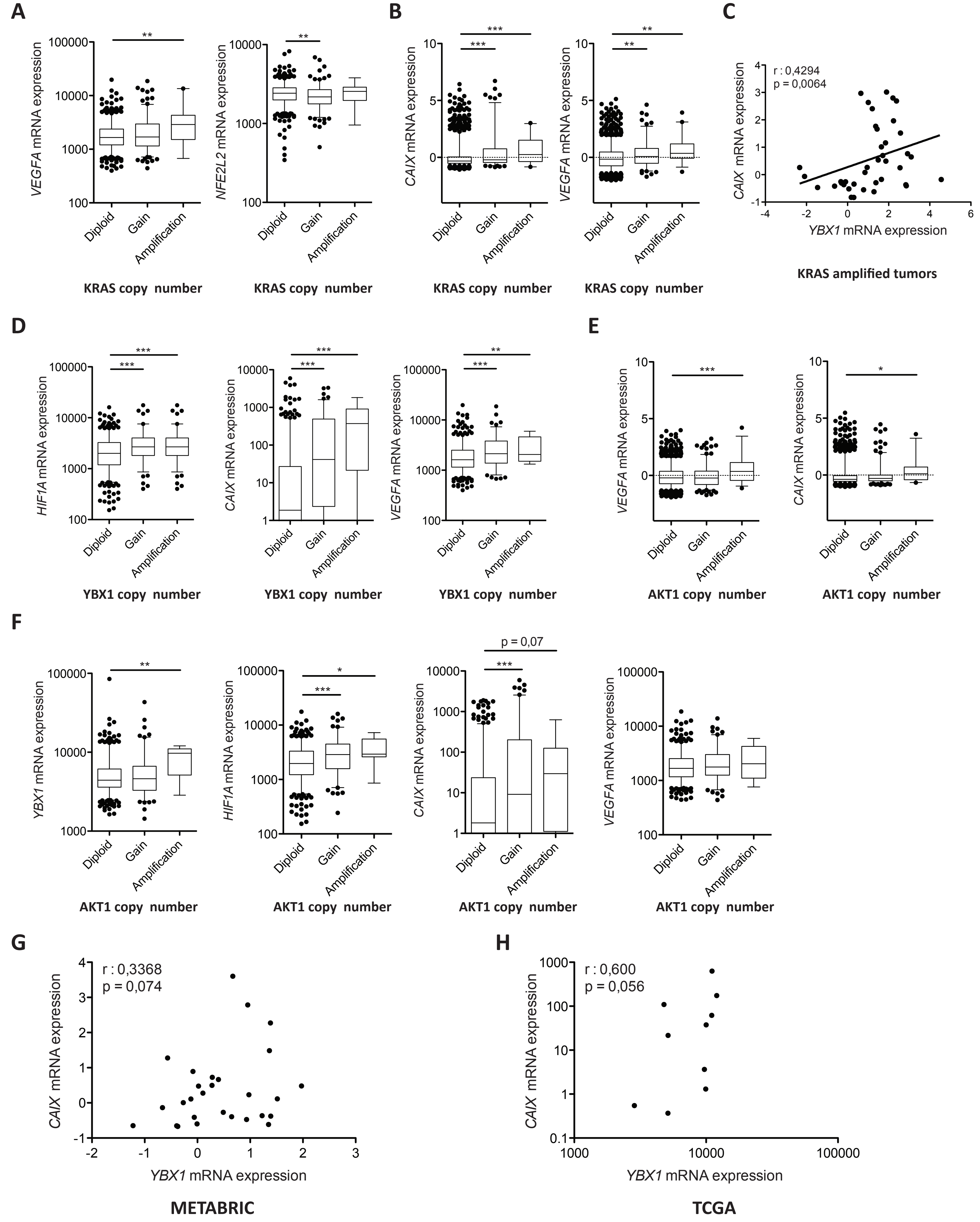

### Appendix Figure S3

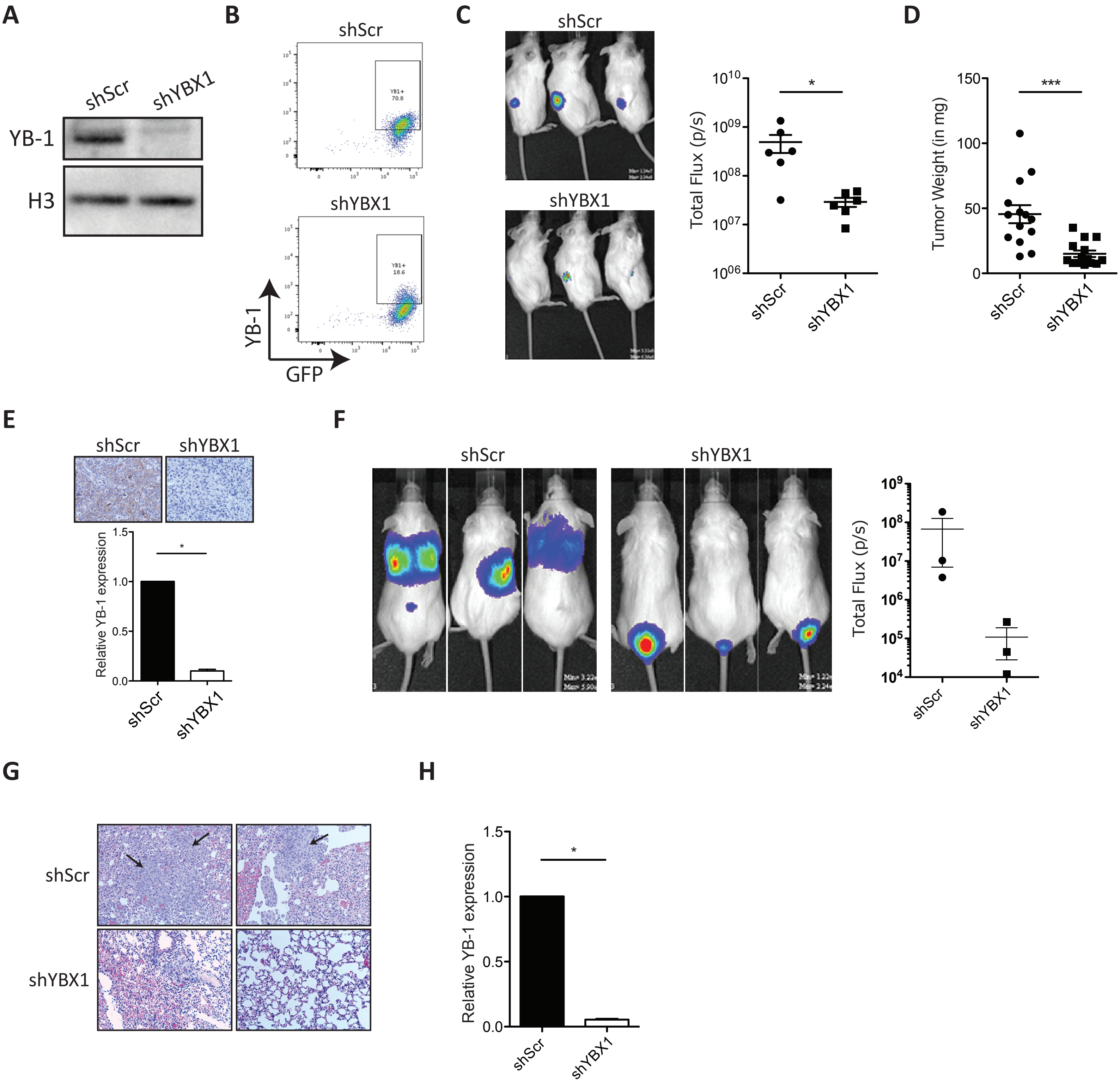

### Appendix Figure S4

**A**

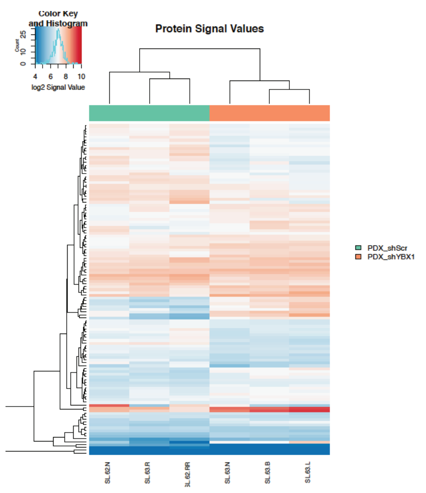


**B**

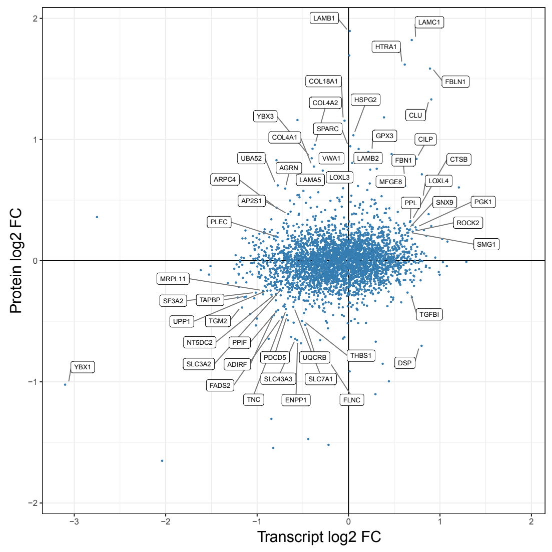


**C**

| | logFC_tumor | logFC_cell | logFC_protein | name |
|---|---|---|---|---|
| CLU | 0,686204948 | 0,561829289 | 1,13 | clusterin |
| AQP1 | 0,685713585 | 0,030616296 | -1,19 | aquaporin 1 (Colton blood group) |
| FBN1 | 0,582909057 | 0,316026985 | 0,33 | fibrillin 1 |
| HTRA1 | 0,566998828 | 0,45267254 | 1,41 | HtrA serine peptidase 1 |
| AKR1C3 | 0,506919019 | 0,282867522 | 0,53 | aldo-keto reductase family 1 member C3 |
| CTSB | 0,465610178 | 0,468705607 | 0,34 | cathepsin B |
| KRT7 | 0,428710908 | 0,251378605 | 0,39 | keratin 7 |
| PPL | 0,420717705 | -0,096063579 | 0,31 | periplakin |
| PTER | 0,395473707 | 0,477052255 | 0,35 | phosphotriesterase related |
| FLNB | 0,372621271 | -0,025828427 | 0,27 | filamin B |
| EML4 | 0,366910152 | 0,095128757 | 0,45 | echinoderm microtubule associated protein like 4 |

**D**

| | logFC_tumor | logFC_cell | logFC_protein | name |
|---|---|---|---|---|
| SLC7A1 | -0,386736896 | 0,021588221 | -0,36 | solute carrier family 7 member 1 |
| BDH1 | -0,39381943 | 0,167863328 | -0,31 | 3-hydroxybutyrate dehydrogenase 1 |
| KRI1 | -0,397984911 | -0,227047716 | -0,32 | KRI1 homolog |
| SLC3A2 | -0,398601902 | -0,081431865 | -0,29 | solute carrier family 3 member 2 |
| TAPBP | -0,414419639 | -0,025529158 | -0,28 | TAP binding protein |
| DNAH2 | -0,461453218 | -0,107131451 | 1,19 | dynein axonemal heavy chain 2 |
| PPIF | -0,477266682 | -0,024606039 | -0,39 | peptidylprolyl isomerase F |
| BRI3BP | -0,56225646 | 0,371085294 | -0,59 | BRI3 binding protein |
| YBX1 | -2,034849551 | -0,223340873 | -0,99 | Y-box binding protein 1 |

**E**

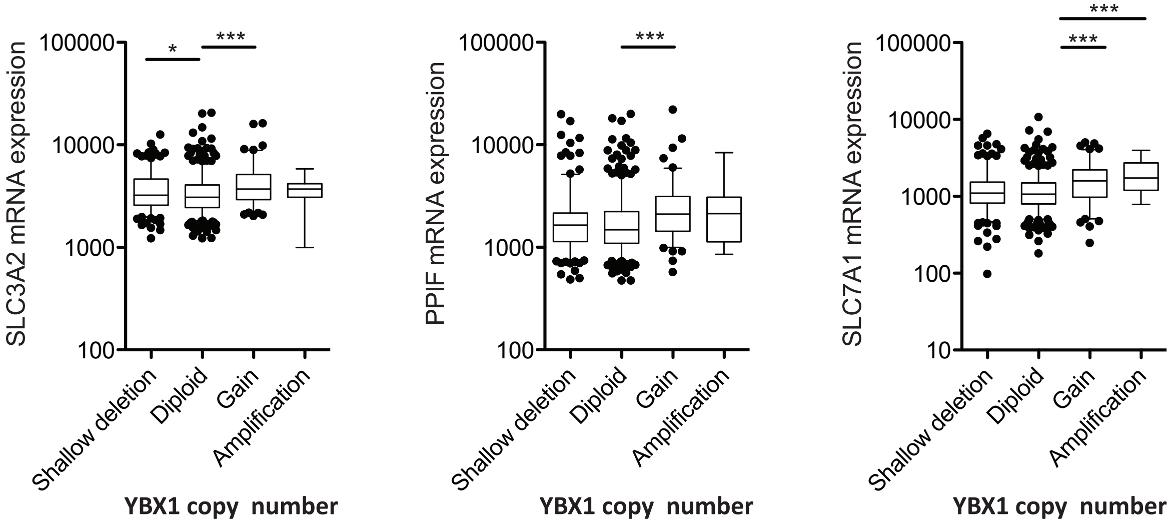

### Appendix Figure S5

A

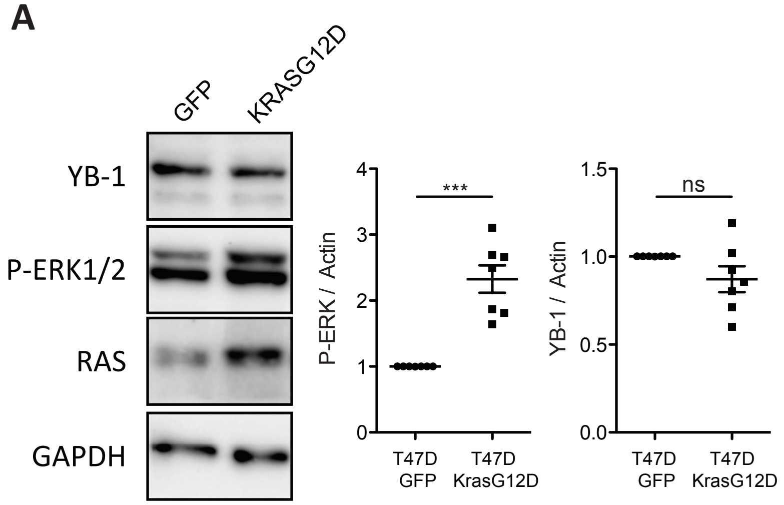


B

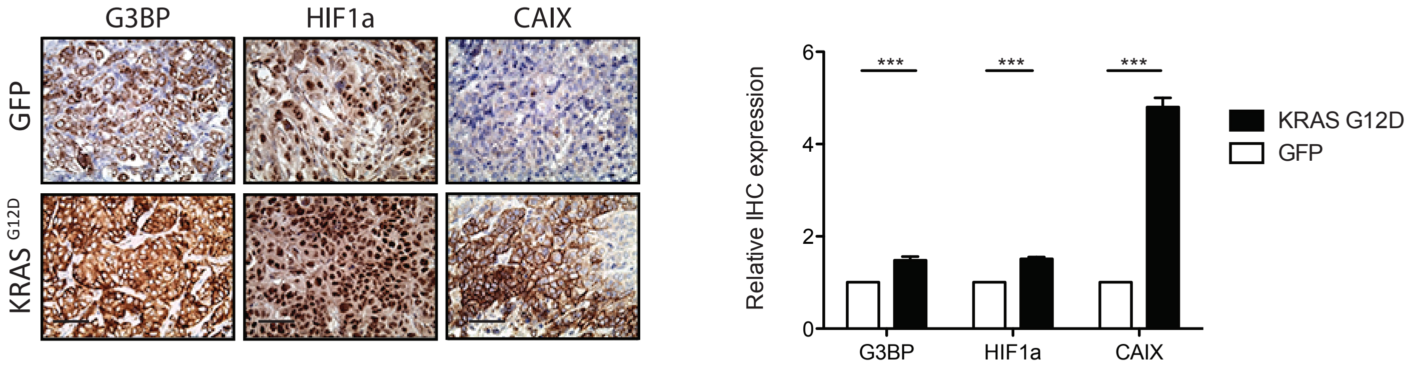


C

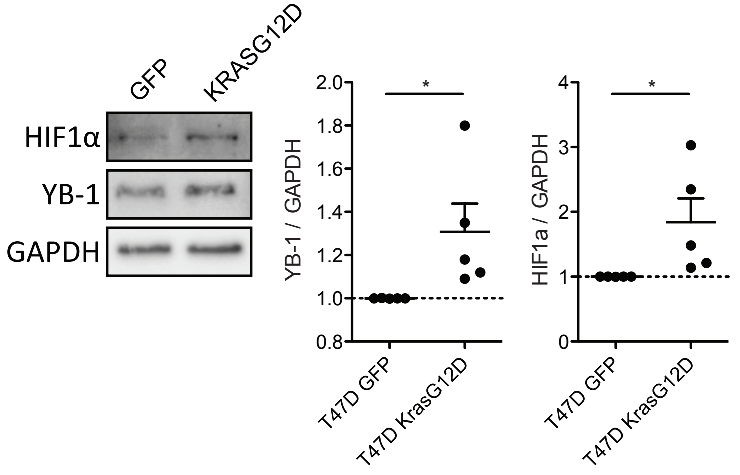

### Appendix Figure S6

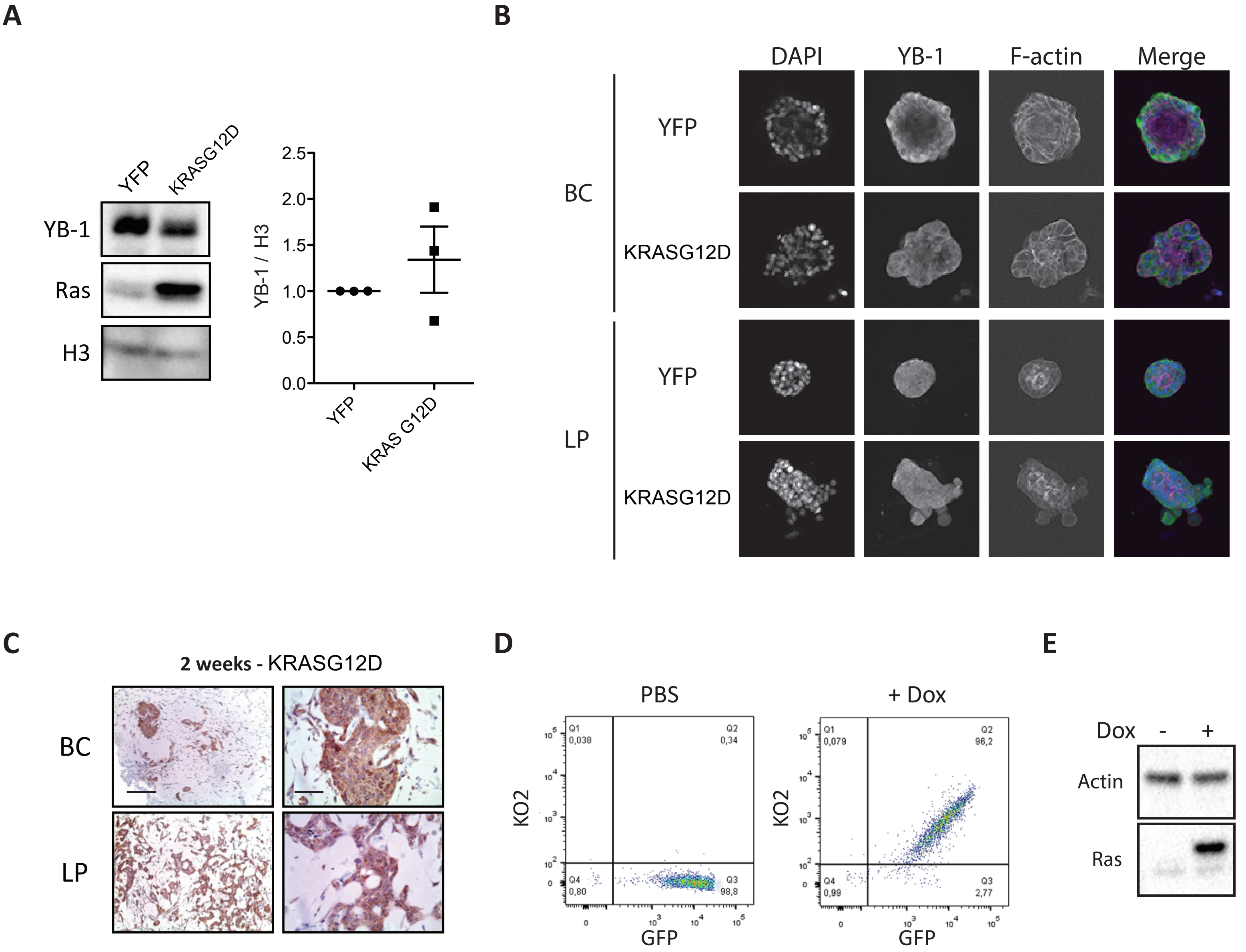

### Appendix Figure S7

**A**

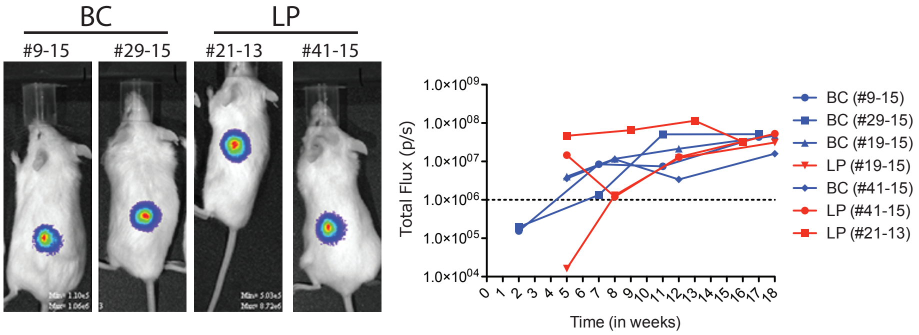


**B**

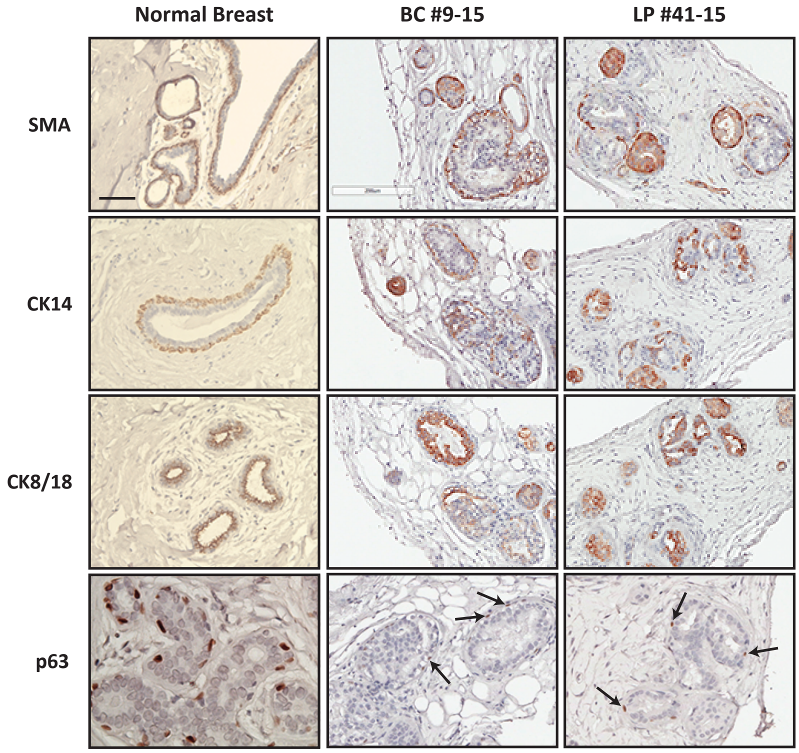


**C**

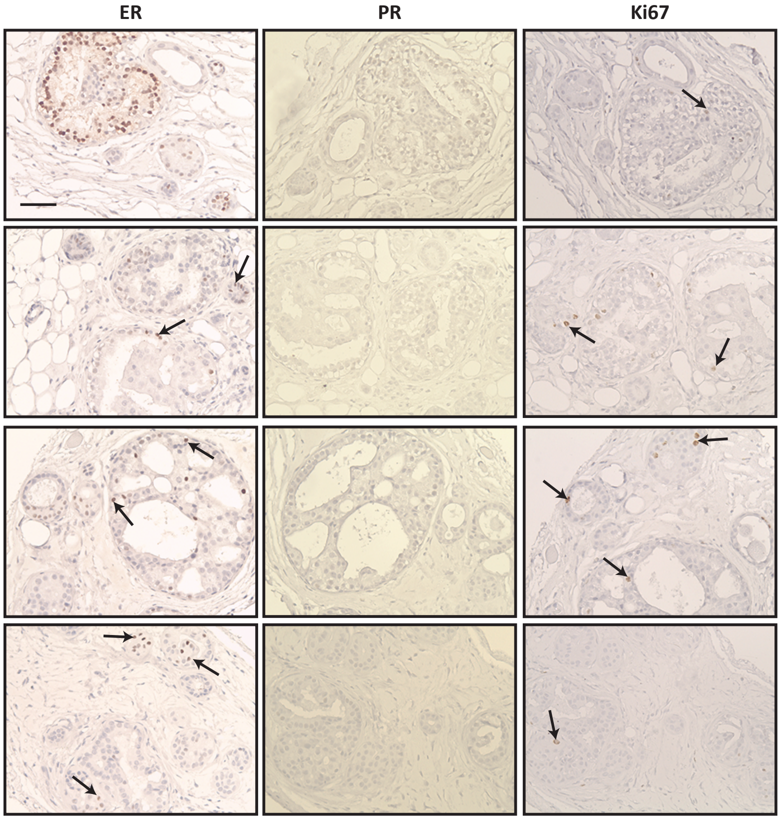

### Appendix Figure S8

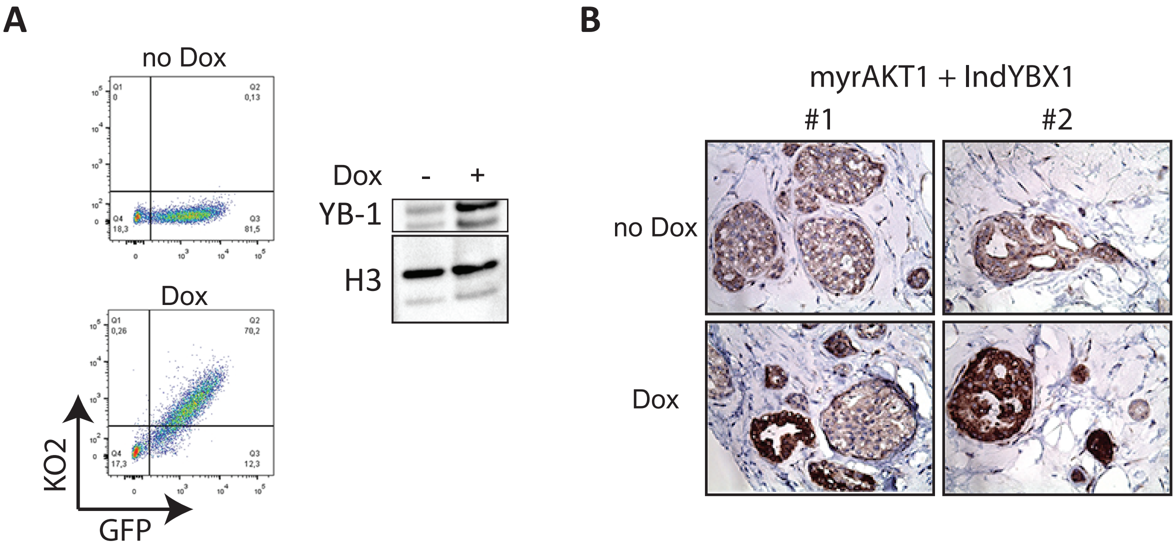

### Supplementary Figure S9

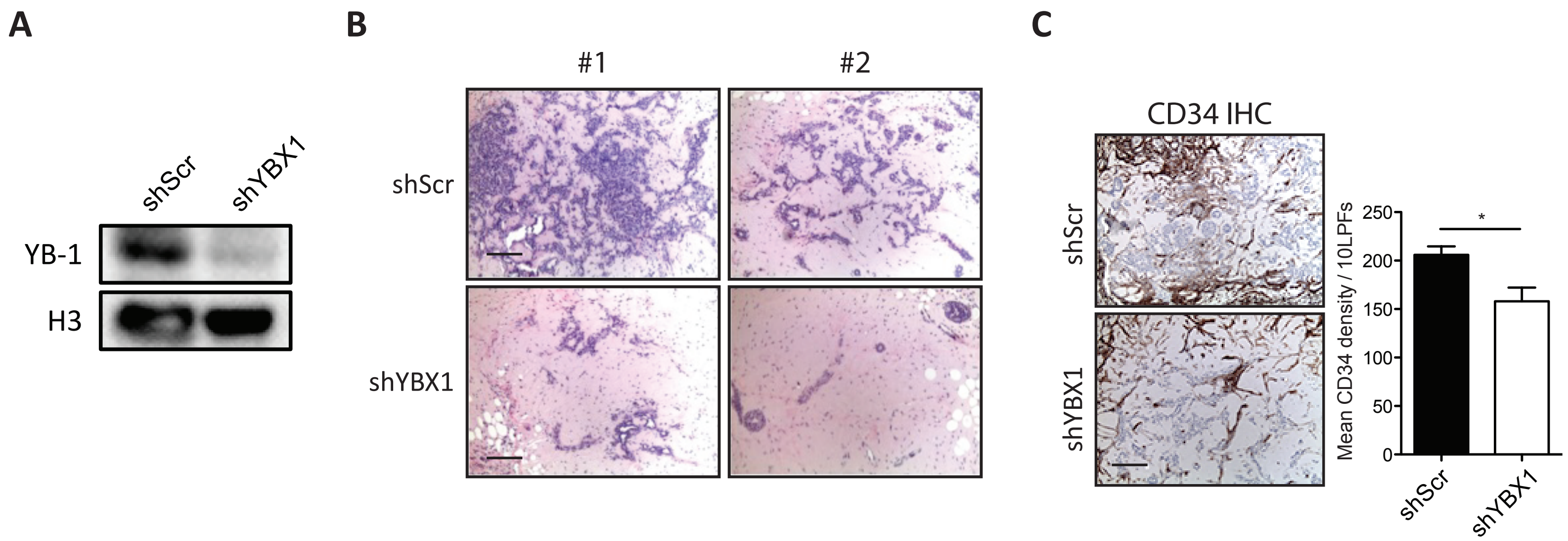
